# Optogenetic reconstitution reveals that Dynein-Dynactin-NuMA clusters generate cortical spindle-pulling forces as a multi-arm ensemble

**DOI:** 10.1101/277202

**Authors:** Masako Okumura, Toyoaki Natsume, Masato T. Kanemaki, Tomomi Kiyomitsu

## Abstract

To position the mitotic spindle within the cell, dynamic plus ends of astral microtubules are pulled by membrane-associated cortical force-generating machinery. However, in contrast to the chromosome-bound kinetochore structure, how the diffusion-prone cortical machinery is organized to generate large spindle-pulling forces remains poorly understood. Here, we develop a light-induced reconstitution system in human cells. We find that induced cortical targeting of NuMA, but not dynein, is sufficient for spindle pulling. This spindle-pulling activity requires dynein-dynactin recruitment/activation by NuMA’s N-terminal long arm, and NuMA’s direct microtubule-binding activities to achieve a multiplicity of microtubule interactions. Importantly, we demonstrate that cortical NuMA assembles specialized focal structures that cluster multiple force-generating modules to generate cooperative spindle-pulling forces. This clustering activity of NuMA is required for spindle positioning, but not for spindle-pole focusing. We propose that cortical <u>D</u>ynein-<u>D</u>ynactin-<u>N</u>uMA (DDN) clusters act as the core force-generating machinery that organizes a multi-arm ensemble reminiscent of the kinetochore.

## Introduction

Forces generated at dynamic plus-ends of microtubules drive directional movement of chromosomes and the mitotic spindle to achieve successful cell division (Inoue & Salmon, 1995). During animal mitosis, dynamic plus-ends of microtubules emanating from the spindle interact with two macro-molecular complexes; kinetochores and the cortical force-generating machinery. Kinetochores consist of more than 100 different proteins assembled on centromeric DNA and surround dynamic microtubule plus-ends using multiple fibril-like microtubule-binding proteins and/or ring-like couplers to harness the energy of microtubule depolymerization for chromosome segregation (Cheeseman, 2014; Dimitrova, Jenni, Valverde, Khin, & Harrison, 2016; McIntosh et al., 2008). In contrast, the cortical force-generating machinery assembles on the plasma membrane and pulls on the dynamic plus-ends of astral microtubules to define spindle position and orientation (Gonczy, 2008; Grill & Hyman, 2005). Spindle positioning determines daughter cell fate by controlling the distribution of polarized cell fate determinants and daughter cell size during both symmetric and asymmetric cell division (di Pietro, Echard, & Morin, 2016; Kiyomitsu, 2015; Morin & Bellaiche, 2011; Williams & Fuchs, 2013). In metaphase human cells, the cortical machinery consists of evolutionary conserved protein complexes, including the cytoplasmic dynein motor, its binding partner dynactin, and the cortically-anchored NuMA-LGN-Gαi complex (Fig. 1A) (Kiyomitsu & Cheeseman, 2012). Prior work has conceptualized that the cortical complex is distributed along the cell cortex and individually pulls on astral microtubules using dynein-based motility and/or by controlling microtubule dynamics (Kiyomitsu & Cheeseman, 2012; Kotak & Gonczy, 2013; Laan et al., 2012). However, compared to the focal kinetochore structure, how this diffusion-prone membrane-associated complex efficiently captures and pulls on dynamic plus-ends of astral microtubules remains poorly understood. Here, we sought to understand the mechanisms of cortical pulling-force generation by reconstituting a minimal functional unit of the cortical force-generating complex in human cells using a light-induced membrane tethering. We found that cortical targeting of NuMA is sufficient to control spindle position, and that NuMA makes multiple, distinct contributions for spindle pulling through its N-terminal dynein recruitment/activation domain, central long coiled-coil, and C-terminal microtubule-binding domains. In addition, we demonstrate that NuMA assembles focal clusters at the mitotic cell cortex that coordinate multiple dynein-based forces with NuMA’s microtubule binding activities. We propose that the cortical <u>D</u>ynein-<u>D</u>ynactin-<u>N</u>uMA clusters (hereafter referred to as the cortical DDN clusters) act as the core spindle-pulling machinery that efficiently captures astral microtubules and generates cooperative pulling forces to position the mitotic spindle.

**Figure 1.**
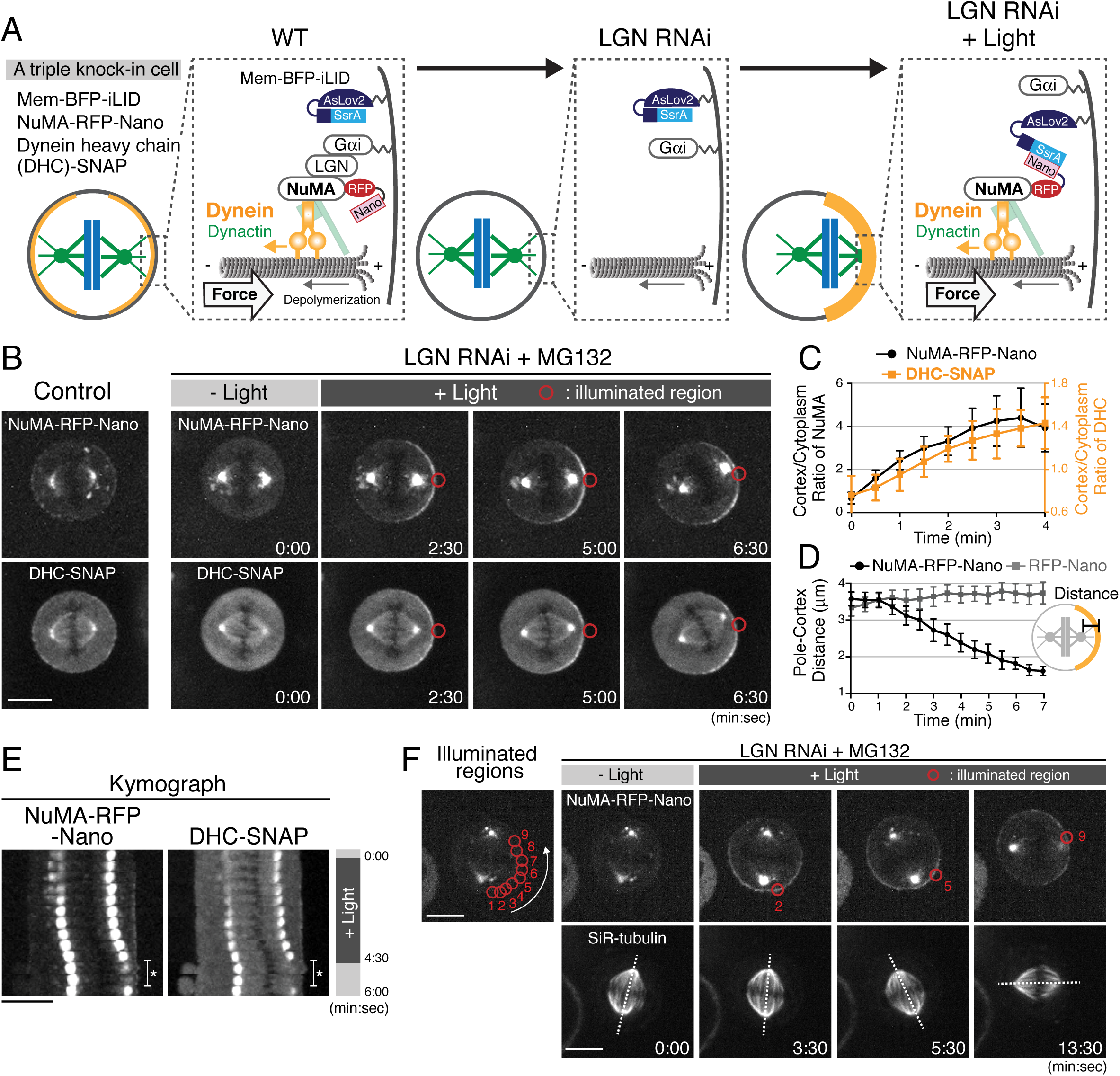
Optogenetic targeting of NuMA to the mitotic cell cortex is sufficient for dynein-dynactin recruitment and spindle pulling. (A) Diagram summarizing cortical complexes in the indicated conditions. (B) Live fluorescent images of NuMA-RFP-Nano (upper) and DHC-SNAP (lower) in control metaphase cells (left), and LGN-depleted cells arrested with MG132. (C) Quantification of cortical NuMA-RFP-Nano and DHC-SNAP signals around the light illuminated region (n=5). Error bars indicate SEM. (D) Quantification of the pole-to-cortex distance (NuMA-RFP-Nano, n=10; RFP-Nano, n=6). Error bars indicate SEM. (E) Kymographs obtained from image sequences in Fig. S2A. Asterisk indicates the duration in which one of the spindle poles moves away from the focal plane. (F) When NuMA-RFP-Nano (upper) was optogenetically repositioned at multiple adjacent cortical regions around the cell membrane by sequential illumination (from 1 to 9), the spindle (lower) rotated about 90° in a directed manner coupled with the changes in cortical NuMA enrichment in 55% (n=11) of cells, but not by repositioning RFP-Nano alone (Fig. S2D, n=6). Dashed lines indicate the spindle axis. Scale bars = 10 μm.

## Results

### Optogenetic targeting of NuMA to the mitotic cell cortex is sufficient for dynein-dynactin recruitment and spindle pulling

To understand the molecular mechanisms that underlie cortical force generation, we sought to reconstitute a minimal functional unit of the cortical force-generating machinery in human cells using a light-induced hetero-dimerization system (iLID) (Guntas et al., 2015). In this system, cytoplasmic RFP-Nano fusion proteins can be targeted to a locally illuminated region of the mitotic cell cortex by interacting with membrane-bound iLID (Fig. 1A, Fig. S1A-B, and Movie S1). Because the N-terminal fragment of NuMA is sufficient to recruit dynein-dynactin to the cell cortex (Kotak, Busso, & Gonczy, 2012), we first sought to manipulate endogenous NuMA. We established triple knock-in cell lines that stably express membrane-targeted BFP-iLID (Mem-BFP-iLID), a NuMA-RFP-Nano fusion (Fig. 1A and Fig. S1C-E), and SNAP-tagged dynein heavy chain (DHC) or the dynactin subunit p150 (Fig. S1F-I). To prevent cortical recruitment of NuMA by the endogenous LGN-Gαi complex, we depleted LGN by RNAi (Fig. 1A middle and 1B t=0:00). We then continuously illuminated the cortical region next to one of spindle poles (indicated by red circles in Figures) with a 488 nm laser to induce NuMA-RFP-Nano targeting. Light illumination induced the asymmetric cortical accumulation of NuMA-RFP-Nano within a few minutes (Fig. 1B-C), which subsequently recruited DHC-SNAP and p150-SNAP (Fig. 1B-C and Fig. S2A-C).

Importantly, following asymmetric NuMA-RFP-Nano targeting, the mitotic spindle was gradually displaced toward the light-illuminated region in 82.4% of cells (n=17, Fig. 1B, D-E, and Movie S2), whereas spindle displacement and cortical dynein recruitment was never observed by targeting RFP-Nano alone (n=6, Fig. 1D and Fig. S2D). Additionally, we found that light-induced repositioning of cortical NuMA is sufficient to drive spindle rotational re-orientation (Fig. 1F and Movie S3). These results indicate that light-induced cortical recruitment of the <u>D</u>ynein-<u>D</u>ynactin-<u>N</u>uMA (DDN) complex is sufficient to generate cortical spindle-pulling forces in human cells.

### Light-induced cortical DDN complex can pull on taxol-stabilized astral microtubules

Cortical pulling forces are supposed to be generated by dynein-based motility on astral microtubules and/or astral microtubule depolymerization coupled with cortical anchorage (Grill & Hyman, 2005). To understand the contributions of astral microtubules to the spindle movement caused by light-induced cortical NuMA, we disrupted or stabilized astral microtubules using the microtubule-targeting drugs, nocodazole or taxol, respectively. In control cells, the metaphase spindle contains visible astral microtubules (Fig. 2A, left) and is displaced following light-induced NuMA-RFP-Nano targeting (Fig. 2B, D-E). In contrast, when astral microtubules were selectively disrupted by treatment with 30 nM nocodazole (Fig. 2A, middle), the spindle was no longer displaced in 56% of cells (n=5/9 cells), and only partially displaced in the remaining 44% of cells (n=4/9) (Fig. 2C-E), despite presence of cortical dynein (Fig. 2C t=5:30). This suggests that astral microtubules are required for spindle pulling by the light-induced cortical DDN complex.

**Figure 2.**
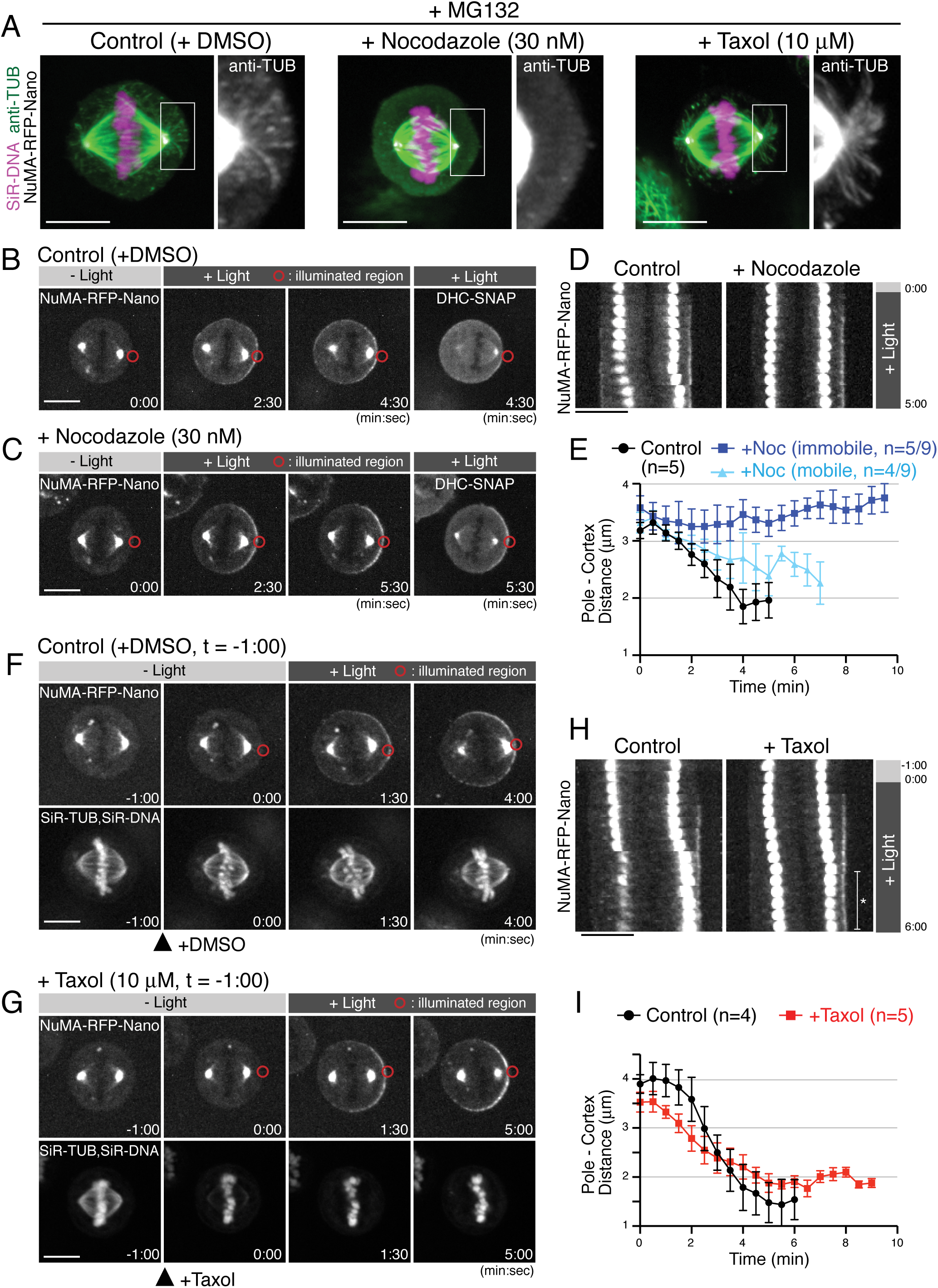
Light-induced cortical NuMA-dynein complex pulls on taxol-stabilized astral microtubules. (A) Fluorescent images of astral microtubules in fixed HCT116 cells treated with drugs as indicated. Cells were arrested at metaphase with MG132 for 1 hr, and DMSO/nocodazole or taxol were then added for 30 min or 1 min, respectively. Images are maximally projected from 15 z-sections acquired using 0.2 μm spacing. (B and C) Live fluorescent images of NuMA-RFP-Nano (upper) and DHC-SNAP (lower) treated with DMSO (B) or nocodazole (C). (D) Kymographs obtained from image sequences in (B) and (C) showing the movement of the spindle at 30-sec intervals. (E) Pole-to-cortex distance for control (black, n=5), and nocodazole-treated cells (blue or light-blue). Blue and light-blue graphs indicate immobile (n=5/9) and partially mobile pools (n=4/9), respectively. Error bars indicate SEM. (F and G) Live fluorescent images of NuMA-RFP-Nano (upper), and SiR-Tubulin and SiR-DNA (Lukinavicius et al., 2015) (lower), treated with DMSO (F) or taxol (G). DMSO and Taxol were added at −1:00, and light illumination began at 0:00, when SiR-tubulin images were selectively abolished by taxol treatment. (H) Kymographs obtained from image sequences in (F) and (G) at 30-sec intervals. In taxol-treated cells, the spindle did not attach to the cell cortex as indicated with an asterisk, likely due to stabilized astral microtubules. (I) Pole-to-cortex distance for control (black, n=4), and taxol-treated cells (red, n=5). Error bars indicate SEM. Scale bars = 10 μm.

Treatment with 10 μM taxol stabilized astral microtubules based on increases in both the length and number of astral microtubules 1 min after addition of taxol (Fig. 2A, right) (Rankin & Wordeman, 2010). Importantly, even in the presence of 10 μM taxol, the spindle was gradually displaced toward the light-illuminated region (Fig. 2H-G t=5:00), although the velocity was slow than that observed in control cells (Fig. 2F-I). In these experiments, we visualized spindle microtubules with 50 nM SiR-tubulin (Lukinavicius et al., 2014), a fluorescent docetaxel derivative, and confirmed the presence of 10 μM taxol by the decrease of SiR-tubulin intensity (Fig. 2G t=0:00), likely due to competition for the same microtubule-binding site. These results suggest that the light-induced cortical DDN complex generates cortical pulling forces by using dynein-based motility on astral microtubules even if microtubule depolymerization is inhibited.

### Light-induced cortical targeting of dynein is not sufficient to pull on the spindle in human cells

A dimerized version of the yeast dynein motor domain is sufficient to position microtubule asters in microfabricated chambers (Laan et al., 2012). To understand the sufficiency of cortical dynein for generating spindle-pulling forces within a human cell, we next directly targeted dynein to the cell cortex (Fig. 3A). Similar to the NuMA-RFP-Nano fusion, endogenously tagged Nano-mCherry-DHC asymmetrically accumulated at the light-illuminated region within several minutes (Fig. 3B and Fig. S3A), and subsequently recruited SNAP-tagged endogenous p150/dynactin to this cortical region (Fig. 3B-C and Fig. S3B). However, endogenous NuMA-SNAP was not recruited to the light illuminated region (Fig. 3D and Fig. S3B), and the spindle was not displaced toward dynein/dynactin-enriched cortex (Fig. 3B right, and 3E) despite the fact that substantial levels of dynein were recruited to the cell cortex (compare Fig. 3C to Fig. 1C). These results suggest that cortical dynein targeting is not sufficient for generating cortical pulling forces in human cells, consistent with recent studies demonstrating that human dynein is auto-inhibited (Torisawa et al., 2014; Zhang et al., 2017) and dynactin and cargo adaptors are required to activate dynein motility (McKenney, Huynh, Tanenbaum, Bhabha, & Vale, 2014; Schlager, Hoang, Urnavicius, Bullock, & Carter, 2014; Zhang et al., 2017).

**Figure 3.**
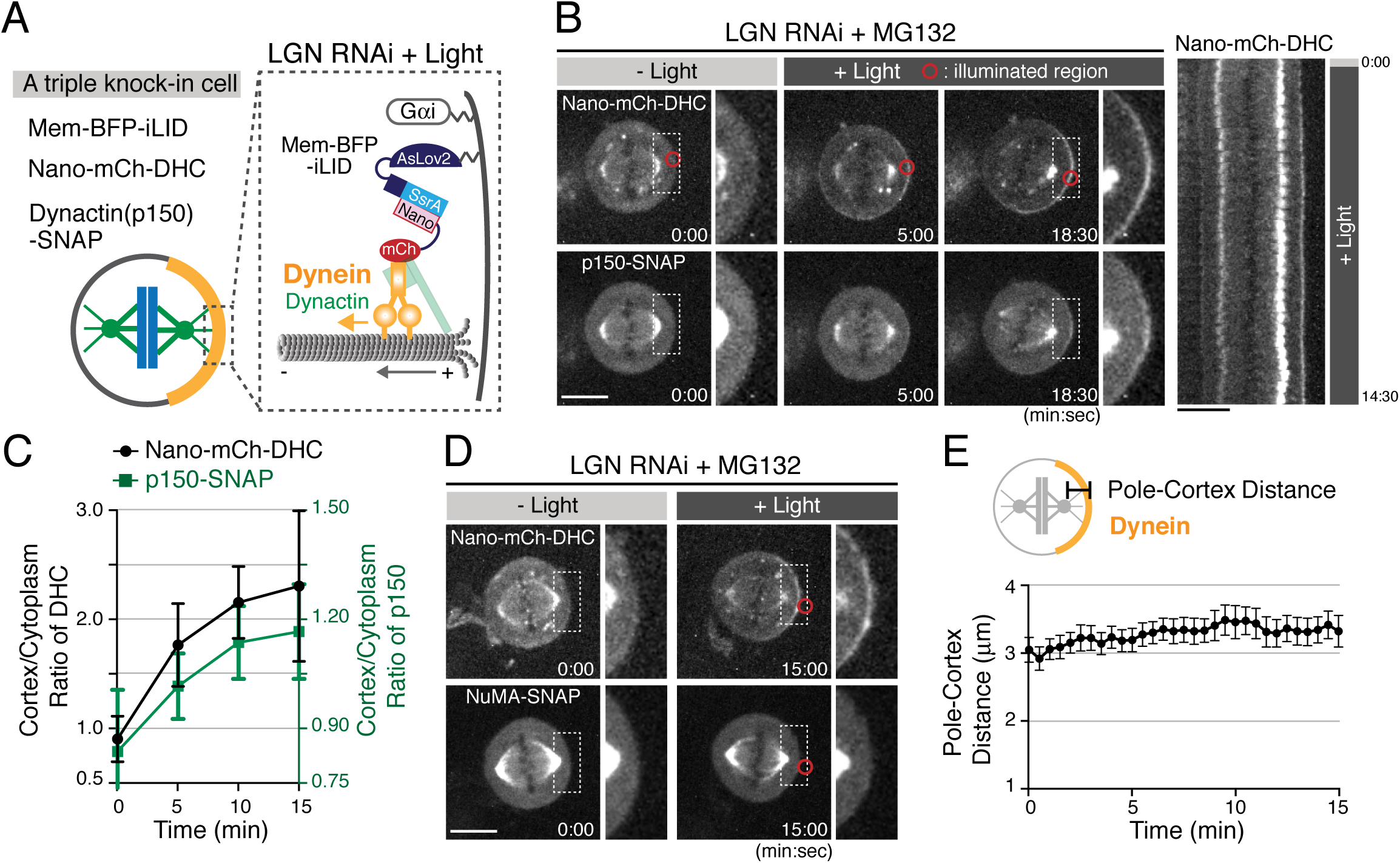
Light-induced cortical dynein targeting is not sufficient to pull on the spindle. (A) Cortical complexes formed by light-induced targeting of Nano-mCherry-DHC. (B) Left: live fluorescent images of Nano-mCherry-DHC (upper) and p150-SNAP (lower). Right: kymograph obtained from image sequences on the left. (C) Quantification of cortical Nano-mCherry-DHC and p150-SNAP signals around the light illuminated region (n=6). Error bars indicate SEM. (D) Live fluorescent images of Nano-mCherry-DHC (upper) and NuMA-SNAP (lower). (E) Measurement of the pole-to-cortex distance (n=10). Error bars indicate SEM. Scale bars = 10 μm.

### A Spindly-like motif in NuMA is required for cortical dynein recruitment, but not sufficient for spindle pulling

The above results suggest that NuMA is required to activate dynein at the cell cortex. Thus, we next sought to define the minimal functional region of NuMA as a dynein adaptor (Fig. 4A). Importantly, our truncation analyses revealed that the NuMA N-terminal region contains a Spindly-like motif sequence (Fig. 4B-E, and Fig. S4A-G) that was recently identified as a conserved binding motif for the pointed-end complex of dynactin in dynein cargo adaptors (Gama et al., 2017). We found that NuMA wild type (WT) fragment (1-705), but not 4A mutant containing alanine mutation in the Spindly-motif (Fig. 4D), recruited dynein to the light-illuminated cortical region (Fig. 4F and Fig. S4H). However, the NuMA (1-705) WT and longer NuMA (1-1700) fragments were unable to fully displace the spindle despite the presence of substantial levels of cortical dynein (Fig. 4B-C, H-I, Fig. S4I-K), whereas ectopically expressed full length NuMA (1-2115 ΔNLS) was able to displace the spindle in ∼40% of cells (Fig. 4G; NLS was deleted to eliminate dimerization with endogenous NuMA.). These results suggest that NuMA recruits dynein-dynactin via its N-terminal Spindly motif, likely to activate dynein’s motility at the mitotic cell cortex similarly to other dynein cargo adaptors (Gama et al., 2017; McKenney et al., 2014; Schlager et al., 2014). However, despite this activation, additional NuMA domains are required to produce cortical spindle-pulling forces.

**Figure 4.**
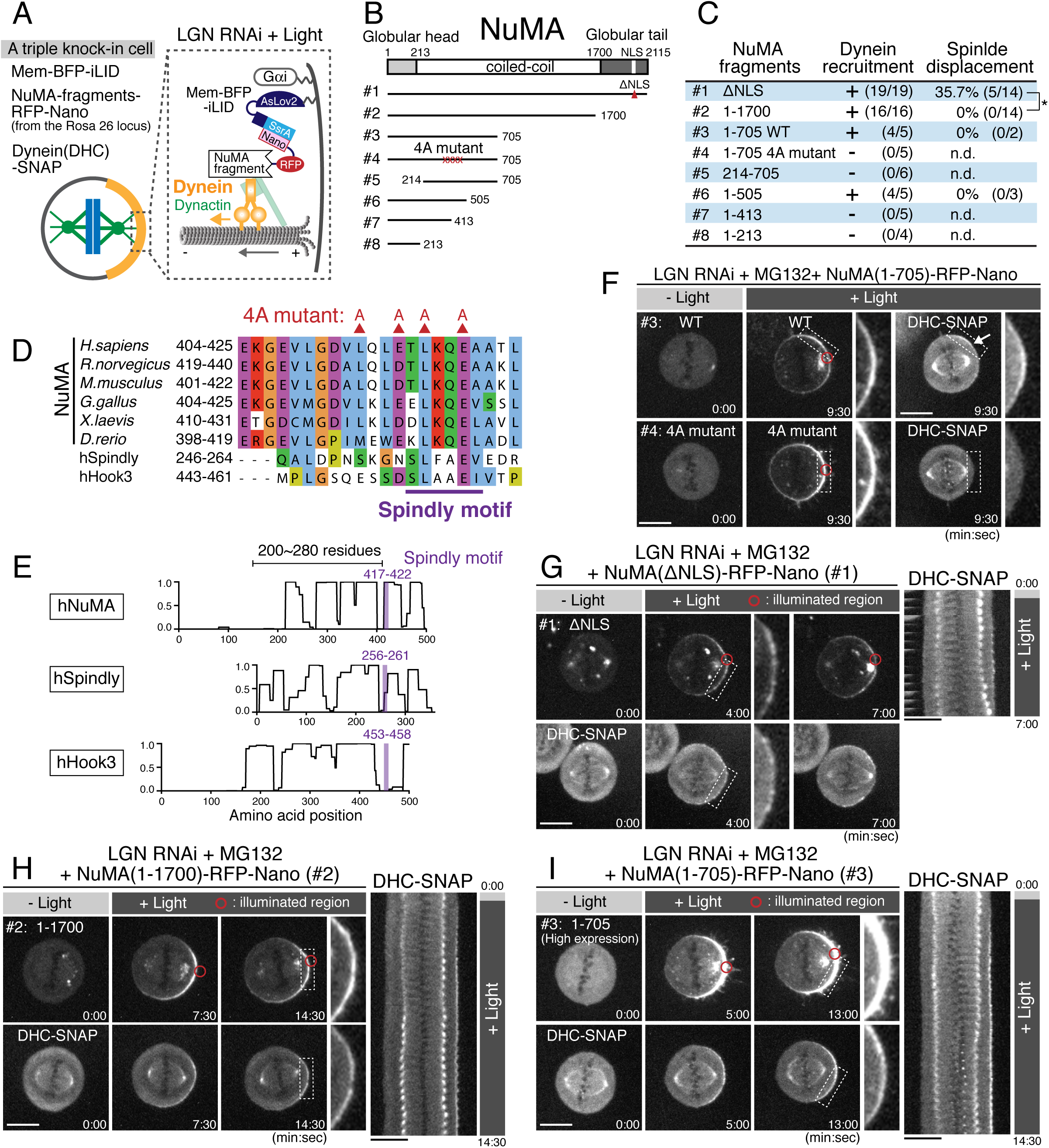
A Spindly-like motif in NuMA is required for cortical dynein recruitment, but not sufficient for spindle pulling. (A) Cortical complexes formed by light-induced targeting of NuMA fragments fused with RFP-Nano. (B) Full-length NuMA and the tested NuMA truncation fragments. Globular domains at N- and C-terminal regions of NuMA are indicated in light-gray and gray, respectively. (C) A summary of the frequency of cortical dynein recruitment and spindle displacement by targeted constructs. Spindle displacement was judged according to the definition of spindle displacement (Fig. S4l). ^*^ indicates significant differences with 99% confidence interval, based on a z-test. (D) Amino acid sequence alignment of the Spindly-motif like region of NuMA proteins in *H. Sapiens* (NP_006176), *R. norvegicus* (NP_001094161), *M. musculus* (NP_598708), *G. gallus* (NP_001177854), *X. laevis* (NP_001081559), *D. rerio* (NP_001316910), and human Spindly (NP_001316568) and Hook3 (NP_115786) aligned by ClustalWS. The conserved L and E substituted by alanine are indicated with red triangles. (E) Lupas coils prediction (window 21). Spindly motif (purple) is commonly located at the C-terminal region of the coiled-coil, with 200∼280 residues. (F) Live fluorescent images of NuMA (1-705) WT (upper) and 4A mutant (lower). DHC-SNAP images are shown to the right. (G-I) Left: live fluorescent images of NuMA constructs (upper) and DHC-SNAP (lower). Right: kymographs obtained from image sequences of DHC-SNAP on the left at 30-sec intervals. Scale bars = 10 μm.

### NuMA’s C-terminal microtubule-binding domains are required for spindle pulling

At kinetochores, a multiplicity of microtubule-binding activities is required to generate cooperative pulling forces (Cheeseman, Chappie, Wilson-Kubalek, & Desai, 2006; Schmidt et al., 2012). Because NuMA’s C-terminal region contains two microtubule-binding domains (MTBD1, and MTBD2) (Fig. 5A and Fig. S5A) (Chang, Huang, Shimamoto, Tsai, & Hsia, 2017; Du, Taylor, Compton, & Macara, 2002; Gallini et al., 2016; Haren & Merdes, 2002), direct binding of NuMA to astral microtubules may generate cooperative forces in parallel with dynein-dynactin recruitment. Consistent with this hypothesis, we found that a Nano fusion with a NuMA (1-1895) fragment, which lacks both microtubule-binding domains, was unable to fully displace the spindle regardless of cortical dynein recruitment (Fig. 5B-C and Fig. S5B). Similarly, NuMA (1-1985), which lacks only the C-terminal microtubule-binding domain (MTBD2), was unable to displace the spindle (Fig. 5B, D and Fig. S5C). In contrast, NuMA Δex24, which lacks exon 24 thus disrupting MTBD1 and an NLS (Fig. 5A) (Gallini et al., 2016; Seldin, Muroyama, & Lechler, 2016; Silk, Holland, & Cleveland, 2009), was able to recruit dynein and displace the spindle similarly to the NuMA-ΔNLS construct (Fig. 5B, E-F). These results indicate that NuMA’s MTBD2 plays critical roles for the ability of the DDN complex to generate spindle-pulling forces.

**Figure 5.**
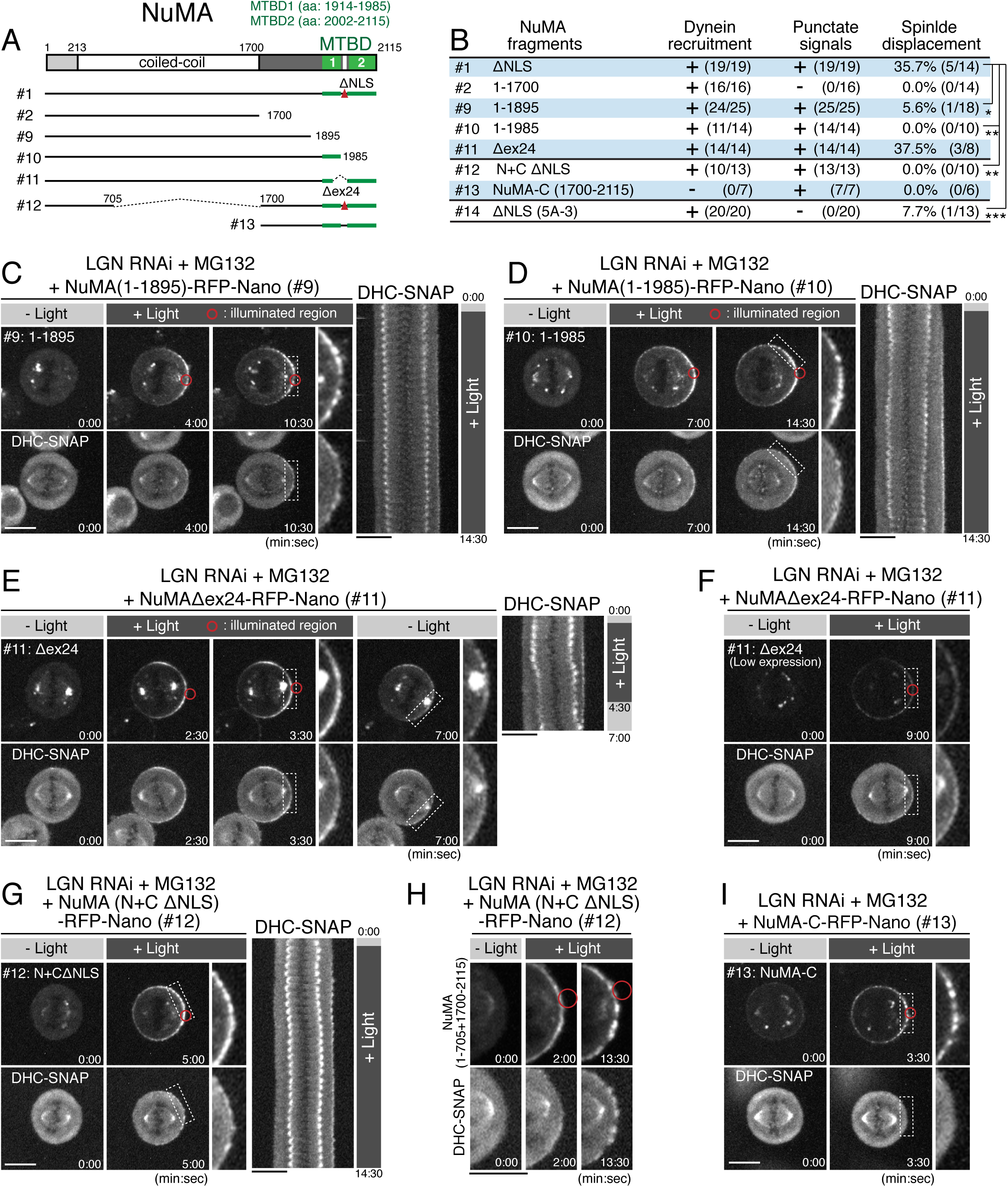
NuMA’s C-terminal microtubule-binding domains and central coiled-coil are required for spindle pulling. (A) Full-length NuMA and the tested NuMA truncation fragments. Microtubule binding domains (MTBDs) are indicated in green. (B) Summary of the frequency of cortical dynein recruitment, dot signal formation and spindle displacement by targeted constructs. ^*^ indicates significant differences, with either 95% (^*^), 99% (^**^), or 90% (^***^) confidence interval, based on a z-test. Spindle displacement was judged according to the definition of spindle displacement (Fig. S4l). (C-E) Left: live fluorescent images of indicated NuMA constructs (upper) and DHC-SNAP (lower). Right: kymographs obtained from image sequences of DHC-SNAP on the left at 30-sec intervals. (F) Live fluorescent images of NuMA Δex24-RFP-Nano (upper) and DHC-SNAP (lower). Expression level of NuMA Δex24-RFP-Nano was lower than that in (E), but the spindle was still displaced. (G) Left: live fluorescent images of NuMA (N+C ΔNLS)-RFP-Nano (upper) and DHC-SNAP (lower). Right: kymographs obtained from image sequences of DHC-SNAP on the left at 30-sec intervals. (H) Enlarged images of NuMA (N+C ΔNLS)-RFP-Nano (upper) and DHC-SNAP (lower) at indicated times. (I) Live fluorescent images of NuMA-C-RFP-Nano (upper) and DHC-SNAP (lower). Scale bars = 10 μm.

### NuMA’s central coiled-coil is required for pulling on the spindle

The work described above defines two important molecular features for cortical force generation: dynein recruitment/activation through the Spindly-like motif and a distinct direct microtubule-binding activity by NuMA. To test whether these features are sufficient to generate cortical pulling forces, we next expressed a fusion construct, NuMA (N+C ΔNLS), that contains both its dynein-recruiting N-terminal and microtubule-binding C-terminal domains, but lacks a ∼1000 aa region of its central coiled-coil (Fig. 5A #12). The NuMA fusion, but not the C-terminal domain (1700-2115) alone (NuMA-C), recruited DHC-SNAP to the light-illuminated region (Fig. 5G-I and Fig. S5D). However, the NuMA (N+C ΔNLS) fusion was unable to fully displace the spindle (Fig. 5G). These results indicate that NuMA’s 200 nm long, central coiled-coil (Harborth, Wang, Gueth-Hallonet, Weber, & Osborn, 1999) also functions with its N-terminal and C-terminal domains to efficiently capture and pull on astral microtubules.

### Identification of a clustering domain on NuMA’s C-terminal region

Our results reveal that NuMA has multiple functional modules for force generation. However, considering the sophisticated kinetochore structure that surrounds a plus-end of microtubule with multiple microtubule-binding proteins (Cheeseman, 2014; Dimitrova et al., 2016), we next sought to define the architecture of the cortical attachment site that is required to efficiently capture and pull on dynamic plus-ends of astral microtubules. Intriguingly, we found that NuMA constructs containing its C-terminal region displayed punctate signals at the cell cortex (e.g. Fig. 5H-I), suggesting that NuMA forms oligomeric structures at the mitotic cell cortex as observed *in vitro* (Harborth et al., 1999). To understand mechanisms of the NuMA’s C-terminal oligomerization/clustering at the mitotic cell cortex, we took advantages of a NuMA-C 3A fragment, which eliminates CDK phosphorylation sites (Compton & Luo, 1995) allowing NuMA to localize to the metaphase cell cortex independently of LGN (Kiyomitsu & Cheeseman, 2013). Similar to the NuMA-C-RFP-Nano (Fig. 5I), GFP-NuMA-C 3A displayed punctate cortical signals (Fig. 6A-B #C1), which was distinct from that of its potential cortical interacting partners - phospholipids, 4.1 proteins and actin cytoskeleton (Kiyomitsu & Cheeseman, 2013; Kotak, Busso, & Gonczy, 2014; Mattagajasingh, Huang, & Benz, 2009; Zheng, Wan, Meixiong, & Du, 2014) – that localize homogenously to the cell cortex (Fig. S6A-C). This suggests that the NuMA C-terminal fragment self-assembles on the membrane.

**Figure 6.**
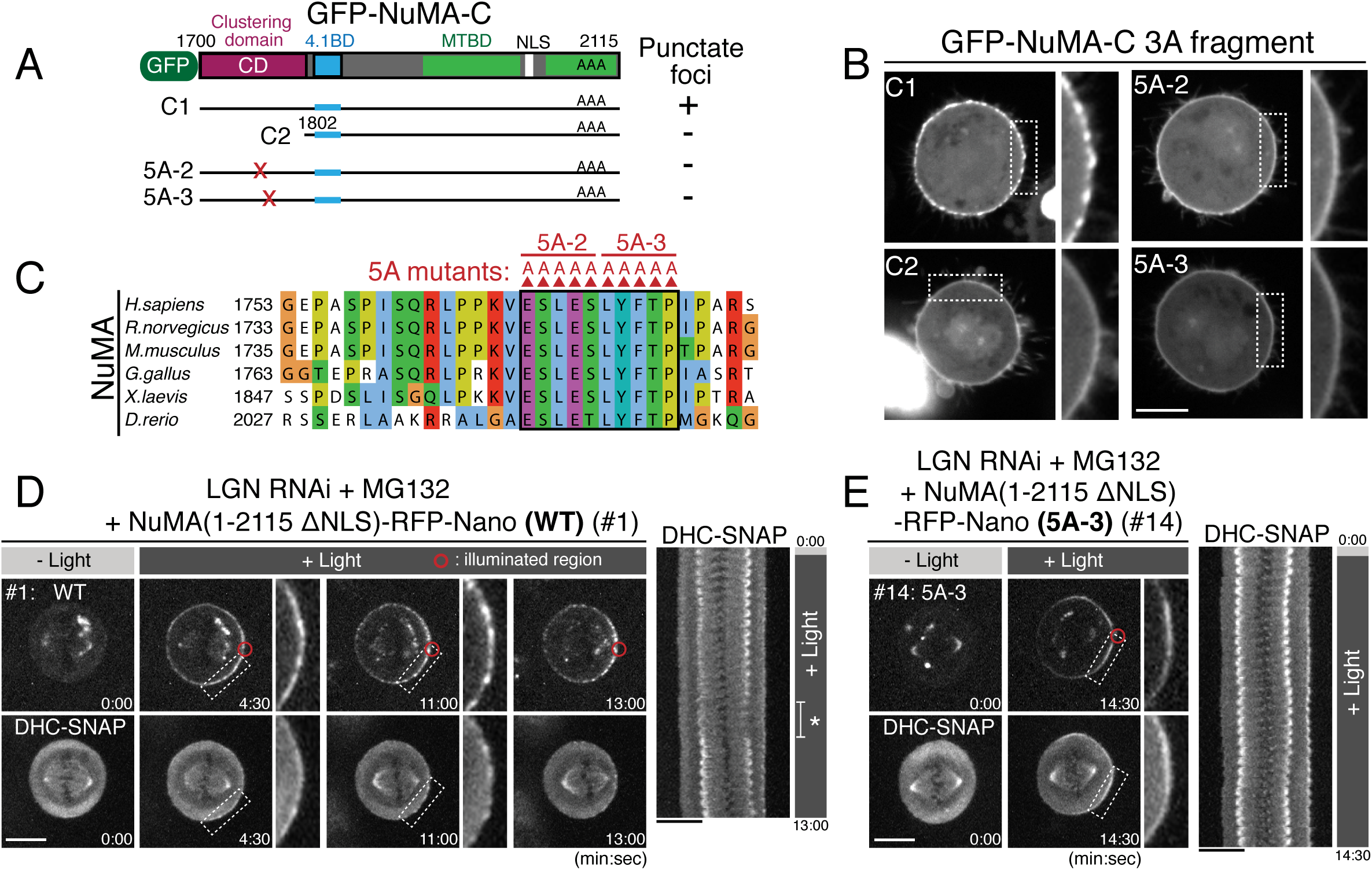
Clustering of the DDN complex by NuMA is critical for spindle pulling. (A) GFP-tagged NuMA C-terminal fragment and the tested NuMA mutant fragments. (B) Live fluorescent images of nocodazole-arrested HeLa cells expressing GFP-tagged NuMA-C 3A fragments. (C) Amino acid sequence alignment of the clustering domain of NuMA proteins aligned by ClustalWS. Accession numbers are indicated in Figure 4D. (D-E) Left: live fluorescent images of indicated NuMA constructs (upper) and DHC-SNAP (lower). Right: kymographs obtained from image sequences of DHC-SNAP on the left. Asterisk in (D) indicates the duration in which one of the spindle poles moves away from the focal plane. Scale bars = 10 μm.

By analyzing different truncations and mutants, we found that a 100 aa region (aa: 1700-1801) of NuMA adjacent to its 4.1 binding domain is required for the formation of punctate foci (Fig. 6A-B, compare #C1 to #C2), and further that a highly conserved 10 amino acid region, E1768-P1777 (Fig. 6C), is necessary for cluster formation (Fig. 6B, see 5A-2 and 5A-3 alanine mutants, and Fig. S6D-F). Consistently, the 1700-1895 region of NuMA is required for the NuMA fragments to display punctate cortical signals (compare Fig. 4H to Fig. 5C, and Fig. S6G). These results suggest an exciting possibility that NuMA assembles a specialized structure to produce large spindle-pulling forces at the cell cortex.

### Clustering by NuMA is required for spindle pulling and positioning, but not for spindle-pole focusing

Above we identified NuMA mutants (5A-2, 5A-3) that are unable to form clusters at the mitotic cell cortex (Fig. 6B-C). To test the functional importance of the novel clustering behavior of NuMA, we next analyzed cortical force generation by full length NuMA wild-type (WT) compared to the 5A-3 mutant using Nano fusions. In cells expressing NuMA (1-2115 ΔNLS)-RFP-Nano (WT), NuMA and DHC-SNAP became gradually detectable as punctate foci (Fig. 6D, 4:30 and 11:00), and the spindle was displaced towards the light-illuminated region (Fig. 6D, 13:00). In contrast, when the NuMA 5A-3 mutant was targeted to the cell cortex, both NuMA 5A-3 mutant and DHC failed to form punctate foci (Fig. 6E, and Fig. S6H), similarly to GFP-NuMA-C 5A-3 (Fig. 6B), and the spindle was not fully displaced (Fig. 5B #14, Fig. 6E and Fig. S6H).

To further probe functional importance of the NuMA’s clustering activity, we next replaced endogenous NuMA with either NuMA WT or the 5A-3 mutant using the auxin-induced degron (AID) system (Fig. 7A) (Natsume, Kiyomitsu, Saga, & Kanemaki, 2016). When endogenous NuMA was degraded, 80% of mitotic cells (n=63) displayed abnormal spindles with unfocused microtubules (Fig. 7B #2 and Fig. S7A-D), consistent with the NuMA KO phenotypes in human hTERT-RPE1 cells (Hueschen, Kenny, Xu, & Dumont, 2017). However, both NuMA WT and the 5A-3 mutant were able to rescue these abnormal spindle phenotypes (Fig. 7B #3 and #4), suggesting that clustering of NuMA is dispensable for microtubule focusing at the spindle poles. In contrast, when endogenous NuMA was replaced with NuMA 5A-3 mutant, the metaphase spindle was tilted and randomly oriented on the x-z plane (26.8 ± 20.7°, n=37, Fig. 7C and Fig. S7E) whereas the spindle in NuMA WT cells was oriented parallel to the substrate (10.7 ± 9.6°, n=34, Fig. 7C) as observed in control metaphase cells (11.5 ± 11.8°, n=41, Fig. 7C). These results suggest that NuMA’s C-terminal clustering is required for proper spindle orientation. We note that the 5A-3 mutation site contains Y1774 (Fig. 6C), which is phosphorylated by ABL1 kinase and contributes to proper spindle orientation (Matsumura et al., 2012). However, treatment with the ABL1 kinase inhibitor Imatinib caused only a mild spindle orientation phenotype (12.3 ± 14.7°, n=27, Fig. 7C), suggesting that the spindle mis-orientation phenotype observed in the 5A-3 mutant is largely attributable to defects in NuMA clustering. Taken together, these results indicate that clustering activity of NuMA is required at the mitotic cell cortex, but not at the spindle poles, for generating cortical pulling forces. Thus, NuMA has a location-dependent structural function that clusters multiple DDN complexes to efficiently capture and pull on dynamic plus ends of astral microtubules.

**Figure 7.**
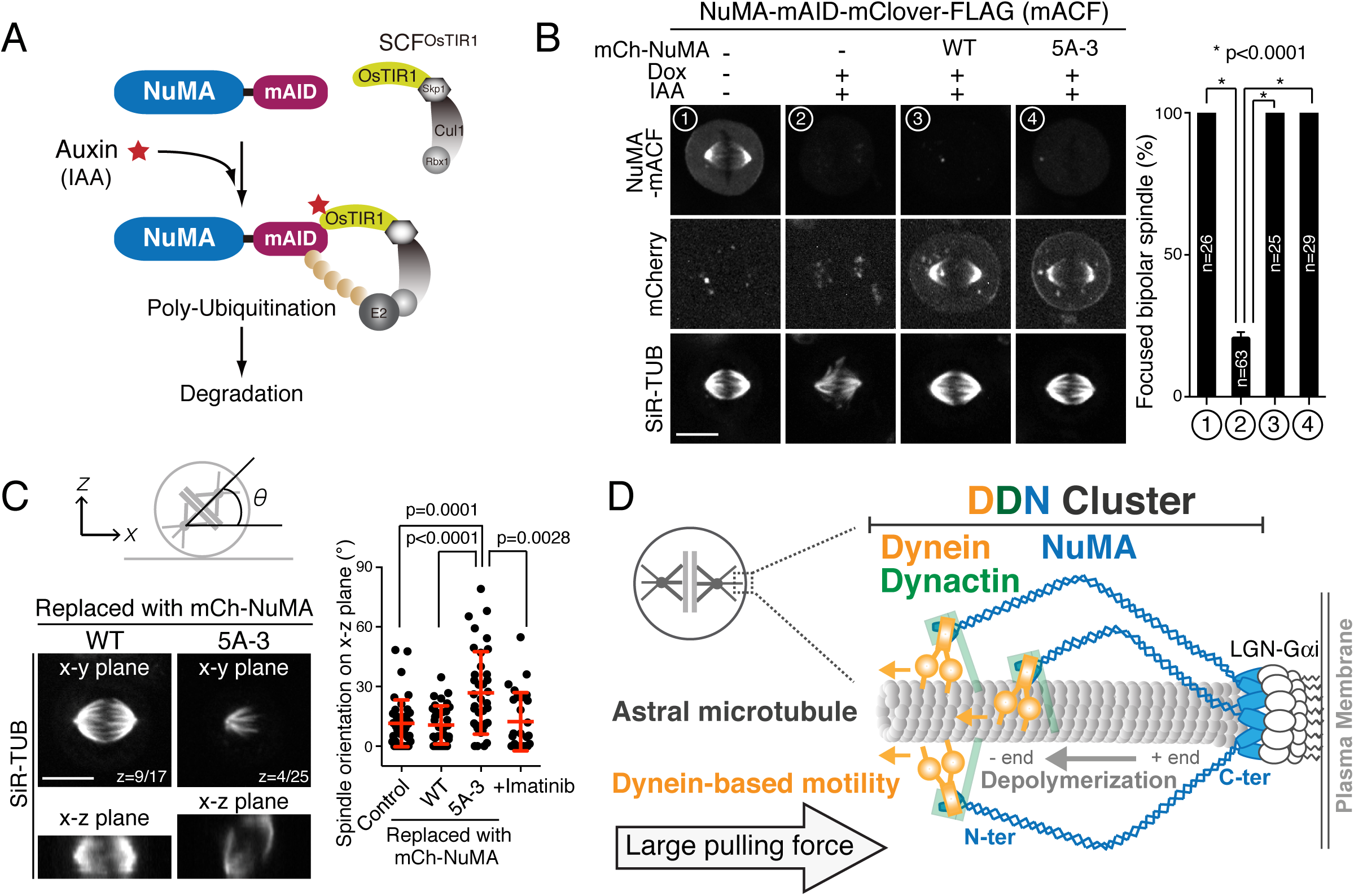
Clustering activity of NuMA is required for spindle positioning, but not for spindle pole focusing. (A) Diagram summarizing auxin inducible degradation (AID) system (Natsume et al., 2016). In the presence of OsTIR1 and auxin (IAA), mAID fusion proteins are rapidly degraded upon poly-ubiquitylation by proteasome. Because RNAi-mediated depletion of NuMA is insufficient to completely deplete NuMA proteins even after 72 hrs (Kiyomitsu & Cheeseman, 2013), we sought to degrade endogenous NuMA using the auxin-induced degron technology. (B) Left: metaphase NuMA-mACF cell lines showing live fluorescent images of NuMA-mACF, mCherry-NuMA WT, or 5A3 mutant, and SiR-tubulin (SiR-TUB) after 24 hrs following the treatment of Dox and IAA. The degradation of endogenous NuMA-mACF was induced by the treatment with Dox and IAA. The expression of mCherry-NuMA WT or 5A-3 was also induced by the Dox treatment. Right: histogram showing frequency of the focused bipolar spindle in each condition. ^*^ indicates statistical significance according to a Student’s *t*-test (P<0.0001). (C) Left: orthogonal views of the metaphase spindle on the x-y (top) and x-z (bottom) plane. In each case, endogenous NuMA was replaced with either mCherry-NuMA WT or 5A-3. Right: scatterplots of the spindle orientation on the x-z plane. Red lines indicate mean ± SD. (D) Model showing multiple-arm capture and pulling of an astral microtubule by the cortical DDN cluster. Scale bars = 10 μm.

## Discussion

### The cortical DDN complex acts as a core functional unit of the cortical force-generating machinery

Here, we applied a light-induced targeting system, iLID (Guntas et al., 2015), for *in cell* reconstitution of the cortical force-generating machinery (e.g. Fig. 1A-B). Our work demonstrates that light-induced targeting of NuMA, but not dynein, is sufficient to control spindle position and orientation in human cells. This is consistent with recent findings that mammalian dynein requires cargo adaptors to activate its motility *in vitro* (McKenney et al., 2014; Schlager et al., 2014; Zhang et al., 2017). In addition, our findings suggest that LGN/Gαi are dispensable for force generation, and instead act as receptors that specify the position of NuMA at the cell membrane. Consistent with this model, LGN-independent pathways that target NuMA to the cell cortex have been reported, such as Dishevelled (Segalen et al., 2010) and phospho-lipids (Zheng et al., 2014). Thus, we propose that the Dynein-Dynactin-NuMA (DDN) complex is a universal core unit that constitutes the cortical force-generating machinery, whereas LGN and other receptors specify the targeting of the DDN complex to the membrane.

### NuMA acts as a force-amplifying platform at the mitotic cell cortex

Our work demonstrates 4 distinct functions for NuMA at the mitotic cell cortex. First, NuMA recruits dynein-dynactin through its N-terminal region. We found that the conserved Spindly-like motif in NuMA is required for dynein recruitment (Fig. 4D-F). NuMA may directly interact with the dynactin pointed-end complex through this Spindly-like motif similarly to other dynein cargo adaptors (Gama et al., 2017), and activate dynein motility at the mitotic cell cortex. Second, the central long coiled-coil of NuMA is required for spindle pulling (Fig. 5G). Purified NuMA displays a long (∼200 nm) rod-shaped structure that shows flexibility with a main flexible-linker region near the middle of central coiled coil (Harborth et al., 1999). Longer flexible arms of NuMA may increase the efficiency of astral microtubule capture by the dynein-dynactin complex, similarly to fibril-like Ndc80 complexes and CENP-E motors at kinetochores (Kim, Heuser, Waterman, & Cleveland, 2008; McIntosh et al., 2008). Third, NuMA contributes to cortical force generation with its own C-terminal microtubule-binding domains (MTBDs) (Fig. 5C), particularly MTBD2 (Fig. 5D). Because this region is also required to prevent hyper-clustering (Fig. S5C right), and is sufficient for cortical localization in anaphase (Fig. S6F C#3, T.K. unpublished observation), this region may play multiple roles for cortical pulling-force generation. Interestingly, a NuMA C-terminal fragment containing MTBD1 (aa: 1811-1985, called NuMA-TIP, Fig. S5A) accumulates at microtubule tips, and remains associated with stalled and/or deploymerizing microtubules (Seldin et al., 2016). By using its two microtubule-binding domains, NuMA may harness the energy of microtubule depolymerization for pulling on astral microtubules similar to the human Ska1 complex or yeast Dam1 ring complex at kinetochores, both of which track with depolymerizing microtubules (Schmidt et al., 2012; Westermann et al., 2006).

Finally, we demonstrate that NuMA generates large pulling forces by clustering the DDN complexes through its C-terminal clustering domain (Fig. 6C-E), similar to lipid microdomains on phagosomes that achieve cooperative force generation of dynein (Rai et al., 2016). Previous studies demonstrated that the 1700-2003 region of NuMA is required for oligomerization *in vitro* (Harborth et al., 1999). We found that 1700-1895 of NuMA is necessary for NuMA fragments to form clusters at the mitotic cell cortex (Fig. 4H and Fig.5C). Because this 1700-1895 fragment itself localizes to the cytoplasm, and showed no punctate signals (Fig. S6D, S6G #C6), the clustering activity of this region may be enhanced by its recruitment and concentration at membranes, as observed for CRY2 clusters (Che, Duan, Zhang, & Cui, 2015). Consistently, NuMA’s clustering function is required for spindle pulling at the cell cortex (Fig. 6E and Fig. 7C), but not for microtubule focusing at spindle poles (Fig. 7B).

### Mechanisms of astral-microtubule capture and pulling by the cortical DDN clusters

Our live-cell imaging revealed that DDN clusters gradually assemble at the cell cortex and then displace the spindle (Fig. 5H, and 6D). Based on the results obtained in this study, we propose a multiple-arm capture model of astral microtubules by the DDN clusters (Fig. 7D and Fig. S7F). Following nuclear envelope break down, cytoplasmic NuMA and DDN complexes are recruited to the mitotic cell cortex by binding to the LGN/Gαi complex, and then assemble DDN clusters on the cell cortex via the NuMA C-terminal domain. *In vitro*, up to 10-12 NuMA dimers self-assemble and form ring-like structures with an average diameter of 48 ± 8 nm (Harborth et al., 1999) (Fig. S7F), which are similar to those of the central hub of yeast kinetochores (37 ± 3 nm) (Gonen et al., 2012), and of the Dam1 ring complex (about 50 nm) which encircles a single kinetochore microtubule (Miranda, De Wulf, Sorger, & Harrison, 2005; Westermann et al., 2005). Given that the NuMA MTBD interacts with depolymerizing microtubules (Seldin et al., 2016), and dynein-dynactin moves along the lattice of microtubules, it is tempting to speculate that the DDN cluster encircles the plus tip of a single astral microtubule with NuMA’s MTBDs, and holds the lateral wall of the astral microtubule with multiple dynein/dynactin-containing arms (Fig. 7D and Fig. S7F). This multiple-arm capture by the DDN cluster leads to larger cooperative pulling forces by increasing the number of both dynein-dynactin containing modules and NuMA’s microtubule binding per an astral microtubule. Additionally, this clustering may contribute to force generation by increasing both the stability of the DDN complex at the membrane, and the frequency for dynein-dynactin to capture or re-bind to astral microtubules. To produce pulling forces at dynamic plus-ends of microtubules, the cortical force-generating machinery appears to develop multiple molecular and structural features analogous to the kinetochore (Cheeseman, 2014; Dimitrova et al., 2016).

In conclusion, our optogenetic reconstitution and AID-mediated replacement reveal that the cortical DDN cluster acts as a core spindle-pulling machinery in human cells. Analyzing the structure and regulation of the DDN cluster will provide further information to understand the basis of spindle positioning in both symmetric and asymmetric cell division, and the general principles for microtubule plus-end capture and pulling.

## Acknowledgments

We thank I. M. Cheeseman and G. Goshima for advice and critical reading of the manuscript, R. Inaba, M. Nishina, K. Murase, and Y. Tsukada for technical assistance, T. Nishiyama and A. Sasaki for reagents, and PRESTO members for discussion. This work was supported by grants from PRESTO program (JPMJPR13A3) of the Japan Science and Technology agency (JST), a Career Development Award of the Human Frontier Science Program, KAKENHI (16K14721, 17H05002) of the Japan Society for Promotion of Science (JSPS), Collaborative Research Program (2014-B, 2015-A1, 2016-A1) of the National Institute of Genetics (NIG), the Uehara Memorial Foundation, the Nakajima Foundation, and the Naito Foundation.

## Author contributions

Conceptualization, T.K.; Investigation, T.K. and M.O.; Formal analysis, T.K., Methodology, T.K., T.N., and M.K; Writing, T.K.; Supervision, T.K.; Funding Acquisition, T.K.

## Materials and methods

- Key resources table

- Plasmid Construction Plasmids for CRISPR/Cas9-mediated genome editing were constructed according to the protocol described in Natsume et al., (Natsume et al., 2016). To construct CRISPR/Cas9 vectors, pX330-U6-Chimeric_BB-CBh-hSpCas9 (#42230, Addgene, Cambridge, MA) was used (Ran et al., 2013). PAM and 20-bp single guide RNA sequences were selected by the optimized CRISPR design tool (http://crispr.mit.edu) (Table S2). To construct donor plasmids containing homology arms for NuMA (∼500-bp homology arms), p150 (∼200-bp arms) and DHC (N-terminal, ∼500-bp arms), a gene synthesis service (Genewiz, South Plainsfield, NJ) was used. To construct the donor plasmid for DHC (C-terminal), a ∼2,000-bp sequence was amplified by PCR from genomic DNA and then cloned into the pCR2.1-TOPO vector. A BamHI site was introduced at the center of the 2,000-bp fragment to facilitate the subsequent introduction of cassettes encoding tag and selection marker genes. To express Mem-BFP-iLID from the AAVS1 locus, membrane-targeted BFP2 (“Mem” from Neuromodulin; Clontech, Mountain View, CA) was fused to the N-terminus of iLID (#60411, Addgene) with a 53-amino acid (aa) linker derived from pIC194 (Kiyomitsu & Cheeseman, 2012) (#44433, Addgene), and the resulting fusion construct was introduced between the AfeI and HindIII sites in pMK231 (AAVS1 CMV-MCS-Puro, #105924, Addgene). Note that the Venus-iLID-caax construct (#60411, Addgene) was able to recruit RFP-Nano, but not NuMA-RFP-Nano to the membrane. To construct the RFP-Nano-NeoR cassette, a tagRFPt-Nano fragment (#60415, Addgene) was introduced between the SacI and MfeI sites in pMK277 (#72793, Addgene). The RFP-Nano-NeoR cassette was excised by BamHI and cloned into the BamHI site in the donor plasmid containing NuMA’s homology arms. A 24-aa linker sequence containing 4× GGGS was introduced between the last codon of NuMA and the first codon of RFP. To construct the Nano-mCherry cassette, the Nano coding sequence was fused to the N-terminal region of mCherry from pIC194 with a 2× GGGS linker. To express Nano-mCherry-DHC, the BSDR sequence from pIC242 (Kiyomitsu & Cheeseman, 2012) (#44432, Addgene) was linked to the Nano-mCherry sequence with a P2A sequence, and the resulting BSDR-P2A-Nano-mCherry cassette, which contained a BamHI site at each end, was inserted into the BamHI site of the donor plasmid for DHC (N-terminal). A 47-aa linker sequence derived from pIC 194 was introduced between the last codon of mCherry and the start codon of DHC. To generate the SNAP-HygroR cassette, the mCherry coding sequence in pMK281 (#72797, Addgene) was replaced with the SNAPf coding sequence (N9186, New England BioLabs, Ipswich, MA) using In-Fusion^®^ cloning (Takara Bio, Ōtsu, Japan). The SNAP-HygroR cassette was excised by BamHI and cloned into the BamHI site of the donor plasmids. To make the DHC donor plasmid containing a SNAP-BSDR cassette, HygroR of the SNAP-HygroR cassette was replaced with BSDR from pIC242 using In-Fusion^®^ cloning. To conditionally express NuMA-RFP-Nano constructs from the Rosa 26 locus, a fragment containing Tet-On 3G, the TRE3GS promoter, and a multiple cloning site (MCS) derived from pMK240 (Tet-On-AAVS1-MCS-PuroR, #105925, Addgene) was introduced into pMK247 (Rosa26-CMV-MCS-HygroR, #105926, Addgene), which contains homology arms for the Rosa 26 locus. An RFP-Nano coding sequence was integrated between MluI and AgeI in the MCS, and NuMA fragments were subsequently inserted into the MluI site. NuMA truncation fragments and mutants were generated by PCR using NuMA cDNA (Compton & Luo, 1995; Kiyomitsu & Cheeseman, 2012) as a template, and the sequences were confirmed by DNA sequencing. These NuMA fragments encode isoform 2 (aa: 1–2101), which lacks a 14-aa region (aa: 1539–1552) in the longer isoform 1. However, the human NuMA constructs presented in the present study conform to isoform 1 (aa: 1–2115; NP_006176) to avoid confusion. To construct mAID-mClover-3×FLAG-NeoR, a 3× FLAG sequence with a GGGS linker was introduced at the C-terminus of mClover of pMK289 (#72827, Addgene) by PCR. To conditionally express mCherry-NuMA WT or the 5A-3 construct from Rosa 26 locus, a fragment containing the TRE3GS promoter and the MCS derived from pMK240 was introduced into pMK247. The mCherry coding sequence derived from pIC 194 was integrated between the MluI and AgeI sites in the MCS, and the NuMA fragments were subsequently inserted Between the SalI and AgeI site.
- Cell Culture and Cell Line Generation HCT116 and HeLa cells were cultured as described previously (Kiyomitsu & Cheeseman, 2012; Natsume et al., 2016; Tungadi, Ito, Kiyomitsu, & Goshima, 2017). Knock-in cell lines were generated according to the procedures described in Natsume et al., (Natsume et al., 2016) with minor modifications. CRISPR/Cas9 and donor plasmids were transfected into the cell lines using Effectene^®^ (Qiagen, Venlo, Netherlands). For drug selection, 1 μg/mL puromycin (Wako Pure Chemical Industries, Osaka, Japan), 800 μg/mL G418 (Roche, Basel, Switzerland), 200 μg/mL hygromycin B (Wako Pure Chemical Industries), and 8 μg/mL blasticidin S hydrochloride (Funakoshi Biotech, Tokyo, Japan) were used. Selection medium was replaced with fresh selection medium 4–5 days after starting selection. After 10–14 days, colonies grown on a 10-cm culture dish were washed once with PBS, picked up with a pipette tip under a microscope (EVOS^®^ XL, Thermo Fisher Scientific, Waltham, MA) located on a clean bench, and subsequently transferred to a 96-well plate containing 50 μL of trypsin-EDTA. After a few minutes, these trypsinized cells were transferred to a 24-well plate containing 500 μL of the selection medium, and then further transferred to a 96-well plate (200 μL per well) for the preparation of genomic DNA. The remaining cells in the 24-well plate were grown and frozen using Bambanker^™^ Direct (Nippon Genetics, Tokyo, Japan). For the preparation of genomic DNA, cells in the 96-well plate were washed once with PBS and then mixed with 60 μL of DirectPCR^®^ lysis solution (Viagen Biotech, Los Angeles, CA) containing 0.5 mg/mL proteinase K (Wako Pure Chemical Industries). The 96-well plate was sealed with an aluminum plate seal and incubated first at 56 °C for 5–6 h, then at 80 °C for 2–3 h in a water bath. To confirm the genomic insertion, PCR was performed using 1–2 μL of the genomic DNA solution and Tks Gflex^™^ DNA polymerase (Takara Bio). The cell lines and primers used in this study are listed in Tables S1 and S3, respectively.
- Microscope System Imaging was performed using spinning-disc confocal microscopy with a 60× 1.40 numerical aperture objective lens (Plan Apo λ, Nikon, Tokyo, Japan). A CSU-W1 confocal unit (Yokogawa Electric Corporation, Tokyo, Japan) with three lasers (488, 561, and 640 nm, Coherent, Santa Clara, CA) and an ORCA-Flash4.0 digital CMOS camera (Hamamatsu Photonics, Hamamatsu City, Japan) were attached to an ECLIPSE Ti-E inverted microscope (Nikon) with a perfect focus system. A stage-top incubator (Tokai Hit, Fujinomiya, Japan) was used to maintain the same conditions used for cell culture (37 °C and 5% CO2). For light illumination, a Mosaic-3 digital mirror device (Andor Technology, Belfast, UK) and a 488-nm laser (Coherent) were used. The microscope and attached devices were controlled using Metamorph (Molecular Devices, Sunnyvale, CA).
- Immunofluorescence and Live Cell Imaging For immunofluorescence in Figure 2A, cells were fixed with PBS containing 3% paraformaldehyde and 2% sucrose for 10 min at room temperature. Fixed cells were permeabilized with 0.5% Triton X-100™ for 5 min on ice, and pretreated with PBS containing 1% BSA for 10 min at room temperature after washing with PBS. Microtubules and DNA were visualized using 1:1000 anti-α-tubulin antibody (DM1A, Sigma-Aldrich, St. Louis, MO) and 1:5000 SiR-DNA (Spirochrome), respectively. Images of multiple z-sections were acquired by spinning-disc confocal microscopy using 0.2-μm spacing and camera binning 1. Maximally projected images from 15 z-sections were generated with Metamorph. For time-lapse imaging of living cells, cells were cultured on glass-bottomed dishes (CELLview™, #627870, Greiner Bio-One, Kremsmünster, Austria) and maintained in a stage-top incubator (Tokai Hit) to maintain the same conditions used for cell culture (37 °C and 5% CO_2_). Three z-section images using 0.5-μm spacing were acquired every 30 s with camera binning 2. Maximally projected z-stack images were shown in figures unless otherwise specified. Microtubules and actin were stained with 50 nM SiR-tubulin and 50 nM SiR-actin (Spirochrome), respectively, for >1 h prior to image acquisition. DNA was stained either 20 nM SiR-DNA (Spirochrome) or 50 ng/mL Hoechst^®^ 33342 (Sigma-Aldrich) for > 1 h before observation. To visualize SNAP-tagged proteins, cells were incubated with 0.1 μM SNAP-Cell^®^ 647 SiR or TMR-STAR (New England BioLabs) for > 2 h, and those chemical probes were removed before observation. For drug treatment, cells were incubated with drugs at the following concentrations and duration: nocodazole, 330 nM (high dose) for 18–24 h and 30 nM (low dose) for 1–4 h; paclitaxel, 10 μM for 1–10 min; cytochalasin D, 1 μM for 1– 10 min; MG132, 20 μM for 1–4 h (Fig. S4B); RO-3306, 9 μM for 20 h; imatinib, 10μM for 24 h (Matsumura et al., 2012); doxycycline hyclate (Dox), 2 μg/mL (Fig. S4B). To express NuMA-RFP-Nano constructs from the Rosa 26 locus in LGN-depleted cells, cells were treated with LGN siRNA (Kiyomitsu & Cheeseman, 2012) and Dox at 24 h and 48 h, respectively, according to the procedure described in Fig S4B. RO-3306 was added at 48 h to cells that were then synchronized at G2 at 68 h. The NuMA-RFP-Nano fusion protein was expressed in most cells, but its expression frequency was reduced in cells that expressed longer NuMA fragments. To compare the intensities of cortically targeted NuMA-Nano fusions, images of NuMA-Nano fusions and DHC-SNAP were acquired using the same parameters (Exposure time: NuMA, 1000 msec; DHC, 500 msec), except for Fig. 1B (NuMA, 1500 msec; DHC, 500 msec). To optimize image brightness, same linear adjustments were applied using Fiji and Photoshop. Supplemental movie files were generated using Metamorph and Fiji. To activate the auxin-inducible degradation of NuMA-mAID-mClover-3FLAG (mACF), cells were treated with 2 μg/mL Dox and 500 μM indoleacetic acid (IAA) for 20–24 h. Cells with undetectable mClover signals were analyzed. A small population of cells showed mClover signals even after being treated with Dox and IAA. For replacement experiments, either mCherry-NuMA WT or the 5A-3 mutant was expressed from the Rosa 26 locus following Dox treatment. This caused the cells to simultaneously express OsTIR1 from the AAVS1 locus to initiate the auxin-inducible degradation of endogenous NuMA-mACF.
- Light-inducible Targeting Except for Figure S1B, HCT116 cells expressing Mem-BFP-iLID and NuMA-Nano fusion proteins were treated with RO-3306 and MG-132 according to the procedure described in Figure S4B to increase the proportion of metaphase-arrested cells. To target Nano fusion proteins at the metaphase cell cortex, cells were illuminated using a Mosaic-3 digital mirror device (Andor Technology) at the indicated regions (circles with a diameter of 3.03 μm) with a 488-nm laser pulse (500 msec exposure, 25 mW). To manually control the frequency of the light pulse and the position of the illuminated region during time-lapse experiments, a custom macro was developed using Metamorph. Using this macro, indicated regions were illuminated ∼10 times with the light pulse during time intervals (30 sec) between image acquisitions. The illuminated position was adjusted to precisely illuminate the cortical region of each cell. In response to the expression level of the Nano fusion proteins, the frequency of the light pulse was reduced to prevent the targeting of Nano fusion proteins throughout the cell cortex. To reposition NuMA-RFP-Nano at the mitotic cell cortex in Fig. 1F, a cortical region adjacent to the spindle axis was illuminated. The light-illuminated region was changed once the spindle started to move but before the spindle was completely attached to the cell cortex. Spindles that rotated by approximately 90° within 15 min were counted.
- Quantification of Cortical Fluorescent Signals and Spindle Displacement Cortical and cytoplasmic fluorescence intensities were determined using Fiji by calculating the mean pixel intensity along three different straight lines (length 3 μm, width 3 pixels) drawn along the cell cortex showing Nano signals or the cytoplasm near the cell cortex but without any aggregations. The background intensity was subtracted from each measurement. The distance from the pole to the cell cortex was measured using Metamorph or Fiji. Line scans for cortical fluorescence intensity were generated using Fiji by calculating the mean pixel intensity along the segmented line (width 3 pixels) drawn along the cell cortex. Kymographs were generated using Photoshop (Adobe Systems, San Jose, CA). Spindle displacement was judged by the definition given in Figure S4I. In addition, cells that satisfied the following conditions were analyzed; 1. NuMA-RFP-Nano fusion proteins were asymmetrically recruited at the light-illuminated region, but not distributed to a whole cell cortex. 2. The cortical intensities of NuMA-Nano fusion proteins were higher than that of NuMA Δex24-RFP-Nano (Fig. 5F). 3. DHC-SNAP was detectable at the light-illuminated region except for the case of the cortical targeting of NuMA-C (#13). 4. The spindle was monitored for >10 min, and not vertically rotated. 5. The bipolar spindle was properly formed without severe membrane blebbing.
- Statistical Analysis To determine the significance of differences between the mean values obtained for two experimental conditions, Student’s *t*-tests (Prism 6; GraphPad Software, La Jolla, CA) or a Z-test for proportions (Allto Consulting, Leeds, UK) was used as indicated in the figure legends.

| REAGENT or RESOURCE | SOURCE | IDENTIFIER |
| --- | --- | --- |
| *Chemicals* |  |  |
| SiR-tubulin | Spirochrome | Cat# SC002 |
| SiR-DNA | Spirochrome | Cat# SC007 |
| SiR-actin | Spirochrome | Cat# SC001 |
| SNAP Cell 647-SiR | New England BioLabs | Cat# S9102S |
| SNAP Cell TMR-star | New England BioLabs | Cat# S9105S |
| Hoechst 33342 | Sigma-Aldrich | Cat# B2261 |
| Nocodazole | Sigma-Aldrich | Cat# M1404 |
| Paclitaxel | Sigma-Aldrich | Cat# T7402 |
| Cytochalasin D | Sigma-Aldrich | Cat# C8273 |
| MG132 | Sigma-Aldrich | Cat# C2211 |
| RO-3306 | Sigma-Aldrich | Cat# SML0569 |
| Imatinib mesylate | Sigma-Aldrich | Cat# SML1027 |
| Puromycin dihydrochloride | Wako Pure Chemical Industries | Cat# 160-23151 |
| G-418 solution | Roche | Cat# 04727894001 |
| Hygromycin B | Wako Pure Chemical Industries | Cat# 084-07681 |
| Blasticidin S hydrochloride | Funakoshi Biotech | Cat# KK-400 |
| Doxycycline hyclate | Sigma-Aldrich | Cat # D9891 |
| 3-Indoleacetic acid (IAA) | Wako Pure Chemical Industries | Cat # 098-00181 |
| DirectPCR® (cell) | Viagen Biotech | Cat #302-C |
| *Antibodies* |  |  |
| Anti-α-tubulin (clone DM1A) | Sigma-Aldrich | Cat# T9026 |
| Rabbit polyclonal anti-NuMA | Abcam | Cat# ab36999 |
| Rabbit polyclonal anti-DHC | Santa Cruz Biotechnology | Cat# sc-9115 |
| Mouse monoclonal anti-p150 | BD Transduction Laboratories | Cat# 610473 |
| Rabbit polyclonal anti-SNAP | New England BioLabs | Cat# P9310S |
| Rabbit polyclonal anti-OsTIR1 | Kanemaki Laboratory (Natsume et al., 2016) | In-house antibody |
| Rabbit polyclonal anti-phospho S10 histone H3 | Abcam | Cat# ab5176-25 |
| Sheep anti-mouse IgG-HRP | GE Healthcare | Cat# NA931 |
| Donkey anti-rabbit IgG-HRP | GE Healthcare | Cat# NA934 |
| *Software and algorithms* |  |  |
| Photoshop CS5, version 12.0 | Adobe Systems | http://www.adobe.com |
| Fiji | (Schindelin et al., 2012) | https://fiji.sc/ |
| Metamorph | Molecular Devices | https://www.moleculardevices.com |
| GraphPad Prism 6, version 6.0c | GraphPad Software | https://www.graphpad.com |
| Excel | Microsoft | https://products.office.com/ |

## Supplemental information

### Supplemental figure legends

**Figure S1:**
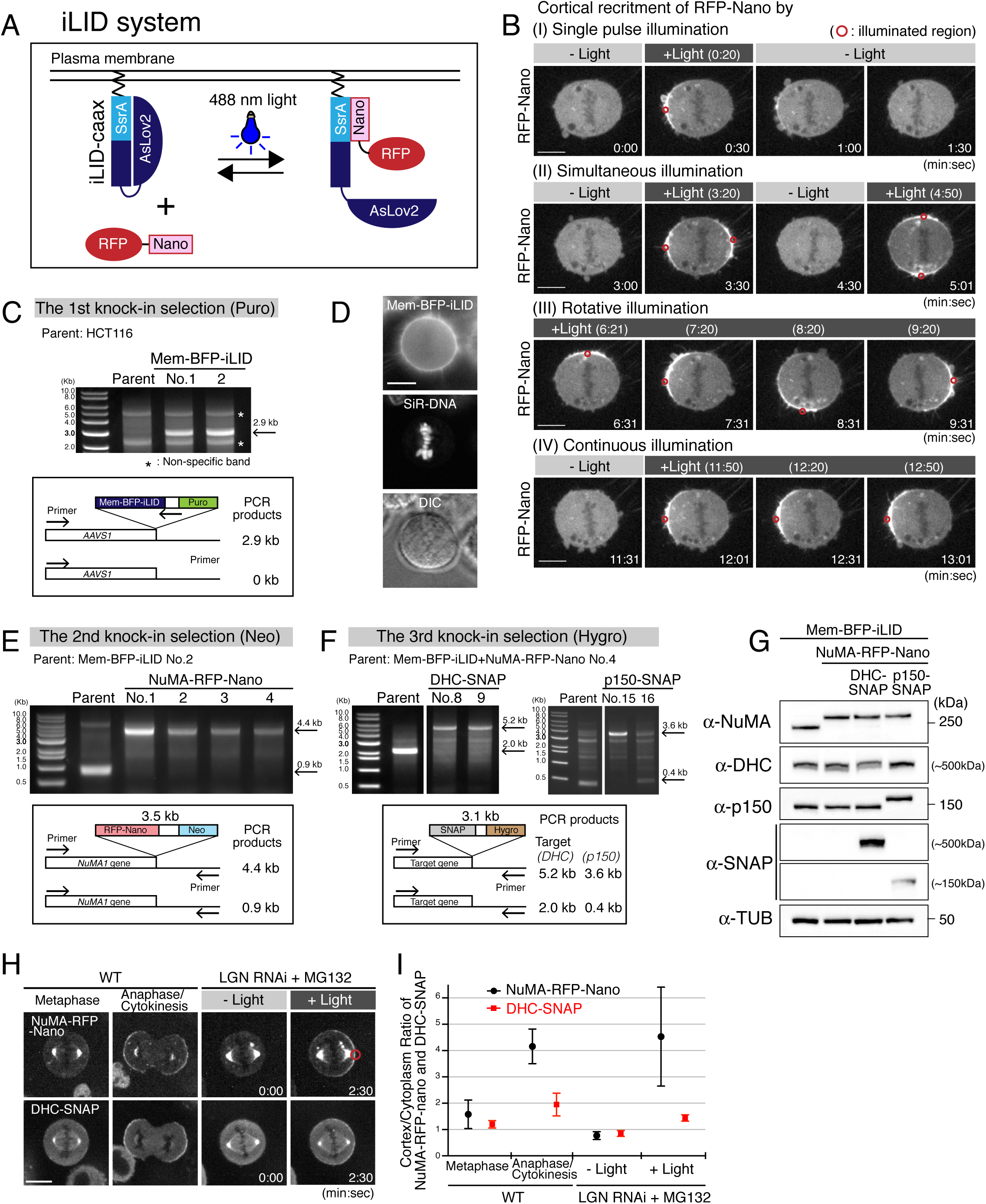
Generation of cell lines for light-induced targeting of endogenous NuMA. (A) Diagram summarizing the iLID system (Guntas et al., 2015). Following blue light illumination, AsLOV2 domain of membrane-targeted iLID induces a conformational change and exposes SsrA peptide, which forms hetero-dimer with Nano-fusions. Upon termination of light illumination, Nano-fusion dissociates from the SsrA with a half-life of <30 sec (Guntas et al., 2015). (B) Time-lapse images of RFP-Nano from a single z-section showing different patterns of cortical RFP-Nano recruitment in response to light illumination. A metaphase HeLa cell transiently expressing RFP-Nano (Addgene #60415) and Venus-iLID-caax (Addgene #60411) was illuminated with a single 488-nm laser pulse (250-msec exposure, 25mW) at the indicated regions (circles with 1.95 μm diameter). See Movie S1. (C) Genomic PCR showing clone genotype after Puromycin (Puro) selection. The clone No.2 was used as a parent in the 2^nd^, and 3^rd^ selection. (D) Live images of Mem-BFP-iLID, DNA, and DIC in the clone No.2 selected in (C). (E) Genomic PCR showing clone genotype after Neomycin (Neo) selection. All clones displayed a single 4.4-kb band, indicating that the RFP-Nano (Neo) cassette was inserted in both *NuMA1* gene loci. The clone No. 4 was used as a parent in the 3^rd^ selection. (F) Genomic PCR showing clone genotype after Hygromycin (Hygro) selection. DHC-SNAP (No. 8, and 9) and p150-SNAP (No. 15) display a single band, as expected, indicating that the SNAP (Hygro) cassette was inserted in both gene loci. The clone DHC-SNAP (No.8) and p150-SNAP (No.15) were used in this study. (G) Western blot probing for anti-NuMA, anti-DHC, anti-p150, anti-SNAP, and anti-α-tubulin (TUB, loading control) showing the bi-allelic insertion of the indicated tags. Protein levels were not significantly affected by tagging with RFP-Nano and SNAP. (H) Live fluorescent images of NuMA-RFP-Nano and DHC-SNAP. NuMA and DHC accumulate at the cell cortex during anaphase (Kiyomitsu & Cheeseman, 2013). (I) Quantification of cortical NuMA-RFP-Nano and DHC-SNAP signals around the polar cell cortex or light-illuminated region (n=5). Error bars indicate SEM. Scale bars = 10 μm.

**Figure S2:**
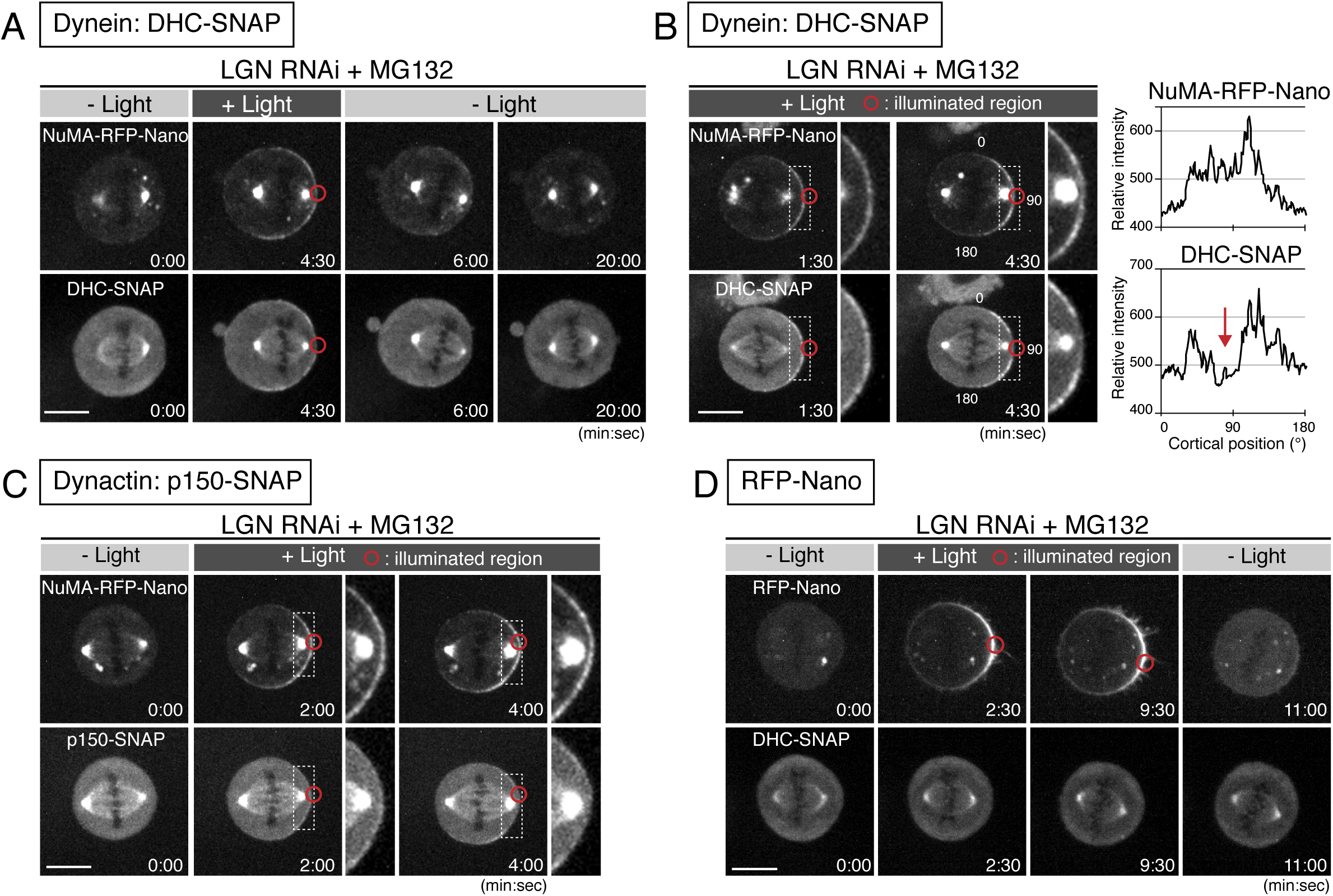
Light-induced cortical targeting of NuMA is sufficient for dynein-dynactin recruitment and spindle pulling. (A) Live fluorescent images of NuMA-RFP-Nano (upper) and DHC-SNAP (lower) in the indicated conditions. Both NuMA-RFP-Nano and DHC-SNAP signals dissociated from the cell cortex following the termination of light illumination (t=6:00), supporting that light-induced NuMA recruits dynein at the cell cortex. Unexpectedly, the displaced spindle gradually returned toward the center of the cell despite the fact that dynein was unable to accumulate at the distal cell cortex to generate opposing cortical pulling forces to center the spindle (t=20:00), suggesting that additional mechanisms exist independently of cortical dynein to center the spindle, and explain why the spindle is roughly positioned in the center of the cell in LGN depleted cells (t=0:00) (Kiyomitsu & Cheeseman, 2012) (B) Left: live fluorescent images of NuMA-RFP-Nano (upper) and DHC-SNAP (lower). Images on the right show a higher magnification of the indicated area. DHC-SNAP signals were initially observed along the cell cortex similarly to NuMA-RFP-Nano (t=1:30), but were selectively diminished from the cell cortex in proximity to the spindle (t=4:30), supporting our model that spindle-pole derived signals negatively regulate the cortical dynein-NuMA interaction in a distance dependent manner (Kiyomitsu & Cheeseman, 2012). Right: line scan showing the relative fluorescence intensity of cortical NuMA-RFP-Nano (upper) and DHC-SNAP (lower) around the cell cortex on the left at 4:30. Arrow indicates a decrease in DHC-SNAP signals near the spindle pole. (C) Live fluorescent images of NuMA-RFP-Nano (upper) and p150-SNAP (lower). Similarly to dynein, p150-SNAP was also recruited to the light illuminated region by NuMA-RFP-Nano (t=2:00), but was subsequently excluded by the spindle proximity (t=4:00). (D) Live fluorescent images of RFP-Nano (upper) and DHC-SNAP (lower) in LGN-depleted cells arrested with MG132. RFP-Nano was expressed from the Rosa 26 locus following Dox treatment (see Fig. S4A-B and S5B). Scale bars = 10 μm.

**Figure S3:**
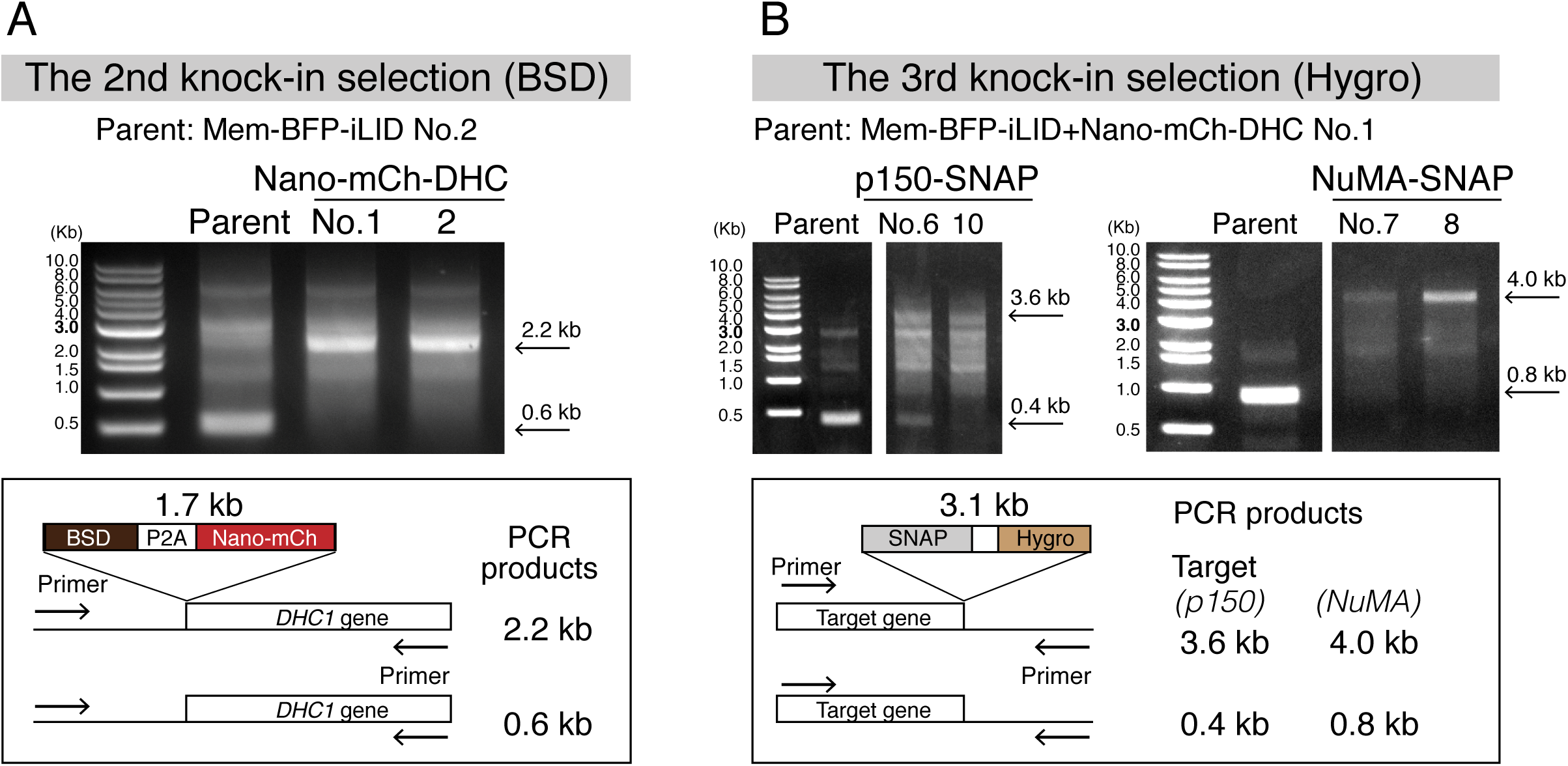
Generation of cell lines for light-induced targeting of endogenous DHC. (A) Genomic PCR showing the genotype of clones after Blasticidin (BSD) selection. Both clones display a single 2.2-kb band, indicating that the Nano-mCherry (BSD) cassette was inserted at both *DHC1* gene loci. The clone No.1 was used as a parent in the 3^rd^ selection. (B) Genomic PCR showing the genotype of clones after Hygromycin (Hygro) selection. P150-SNAP (No.10) and NuMA-SNAP (No.7, 8) display a single band, as expected, indicating that the SNAP (Hygro) cassette was inserted at both gene loci. The clone p150-SNAP (No.10) and NuMA-SNAP (No.8) were used in this study.

**Figure S4:**
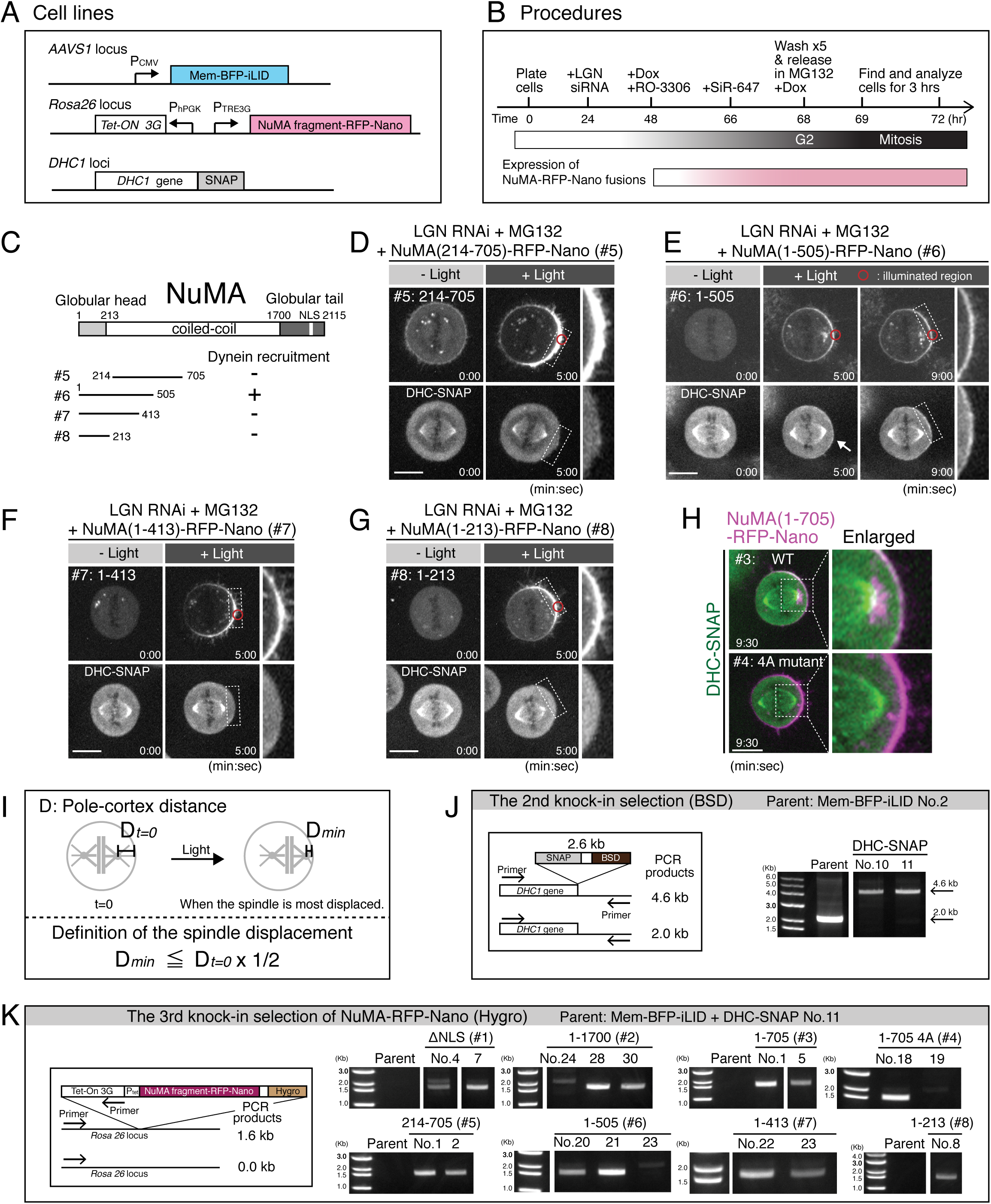
The N-terminal region of NuMA is required for cortical dynein recruitment. (A) A schematic illustration of exogenous gene expression of Mem-BFP-iLID and NuMA-Nano fusions from the AAVS1 and Rosa 26 loci, respectively. Whereas Mem-BFP-iLID is stably expressed, NuMA-Nano fusions are conditionally expressed following the treatment with Doxycycline (Dox). The SNAP-tag was also inserted at the *DHC1* gene loci. (B) Schematic of experimental procedures. LGN siRNA, Dox, and SiR-647 were used to deplete endogenous LGN, to induce expression of NuMA-Nano fusions, and to label endogenous DHC-SNAP, respectively. Cells were treated with RO-3306 and MG132 to synchronize at G2 and to arrest cells at metaphase, respectively. Cells were observed by microscopy 1hr after the release from G2 arrest, and analyzed for 3hrs. (C) Left: the tested NuMA truncation fragments. Right: A summary of the cortical dynein recruitment. (D-G) Live fluorescent images of indicated NuMA constructs (upper) and DHC-SNAP (lower). NuMA (1-505) was sufficient to recruit dynein to the cell cortex, and both its N-terminal globular domain (aa: 1-213) and short coiled-coil region (aa: 414-505) were required for cortical dynein recruitment. Similar to NuMA (1-705) WT, NuMA (1-505), but not other truncated fragments, accumulated around spindle pole adjacent to light-illuminated region following cortical recruitment. (H) Merged images of NuMA(1-705)-RFP-Nano (magenta) and DHC-SNAP (green) from Fig. 4F. NuMA(1-705)-RFP-Nano WT, but not 4A mutant, asymmetrically accumulates around the spindle pole, indicating that NuMA N-terminal fragments containing the Spindly motif are sufficient to recruit dynein-dynactin to the mitotic cell cortex, and likely to activate dynein’s motility, which in turn transports these NuMA fragments toward the spindle pole. (I) The definition of spindle displacement. In cases where D_min_ is smaller than D_t=0_ x 1/2, the spindle was judged as displaced. (J) Genomic PCR showing the genotype of clones after Blasticidin (BSD) selection. Both clones display a single 4.6-kb band, indicating that the SNAP (BSD) cassette was inserted in both *DHC1* gene loci. The clone No.11 was used as a parent in the 3^rd^ selection. (K) Genomic PCR showing the genotype of clones after Hygromycin (Hygro) selection. A 1.6-kb band confirms the insertion of NuMA-RFP-Nano (Hygro) cassettes with different NuMA fragments at the Rosa 26 locus. The following clones were used; ΔNLS (#1: No.7), 1-1700 (#2: No.28), 1-705 (#3: No.5), 1-705 4A (#4: No.18). 214-705 (#5: No.2), 1-505 (#6: No.21), 1-413 (#7: No.23), and 1-213 (#8: No.8), Scale bars = 10 μm.

**Figure S5:**
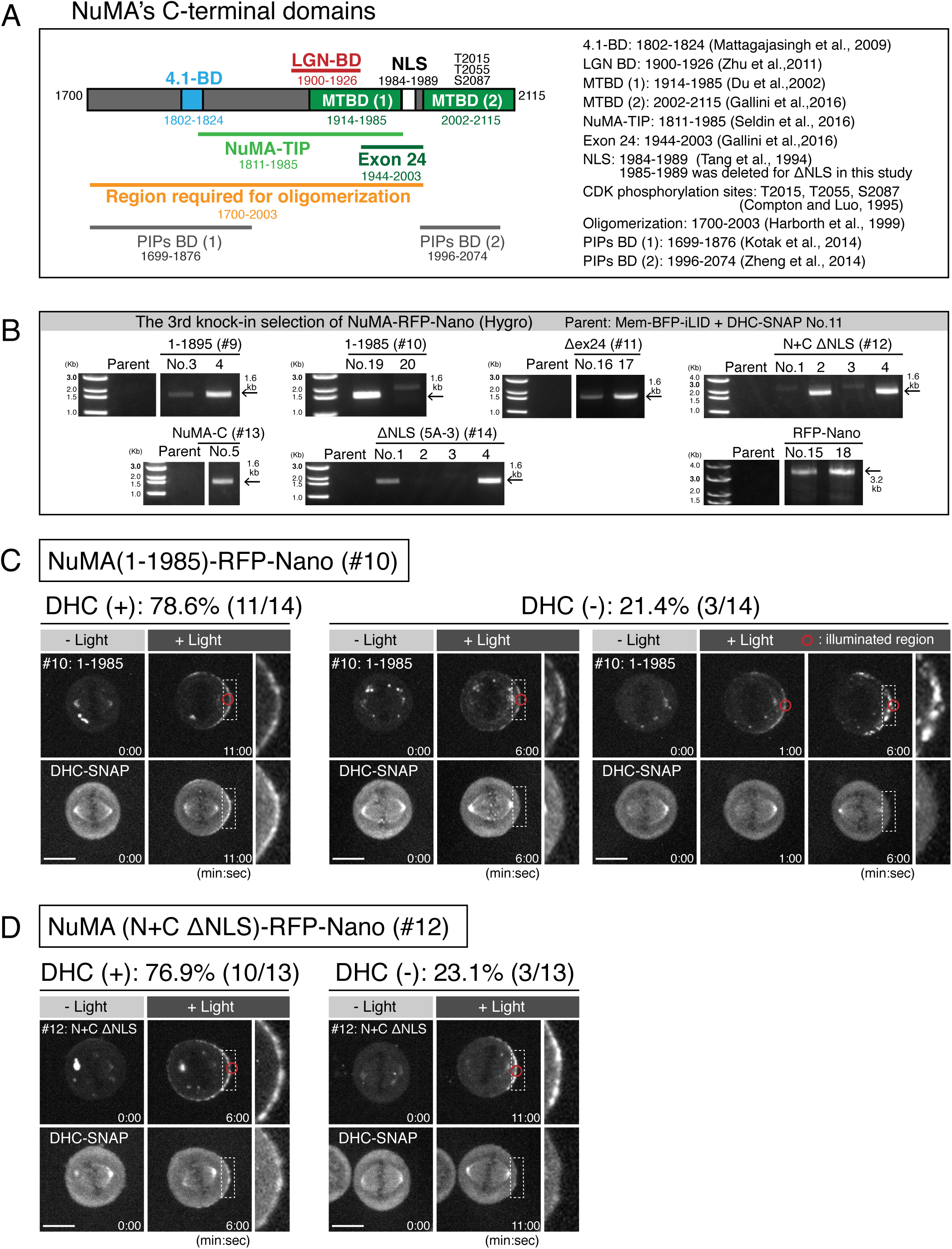
Light-induced targeting of exogenously expressed NuMA fragments lacking C-terminal MTBDs and central coiled-coil. (A) NuMA C-terminal fragment with known domains. 4.1-binding domain (4.1-BD) and microtubule binding domains (MTBDs) are indicated in light-blue and green, respectively. The amino acid numbers of NuMA conform to isoform 1 (aa: 1–2115; NP_006176). LGN binding domain (BD) (Zhu et al., 2011) is indicated in red. NuMA C-terminal fragment containing MTBD1 (aa: 1811-1985, called NuMA-TIP) accumulates at microtubule tips, and remains associated with stalled and/or deploymerizing microtubules (Seldin et al., 2016). (B) Genomic PCR showing the genotype of clones after Hygromycin (Hygro) selection. Arrows indicate a 1.6- or 3.2-kb band, which confirms the insertion of NuMA-RFP-Nano (Hygro) cassettes with different NuMA fragments at the Rosa 26 locus. The following clones were used; 1-1895 (#9: No.4), 1-1985 (#10: No.19), Δex24 (#11: No.17), N+C ΔNLS (#12: No.4), NuMA-C (#13: No.5), ΔNLS(5A-3) (#14: No.4), and RFP-Nano (No.18). (C) Live fluorescent images of NuMA(1-1985)-RFP-Nano (upper) and DHC-SNAP (lower). Whereas DHC was recruited to the cell cortex following light-induced NuMA(1-1985)-RFP-Nano targeting in 78.6% of cells (n=14), DHC was not detectable in the remaining 21.4% of cells, in which NuMA(1-1985)-RFP-Nano fusion apparently aggregates at the cell cortex, suggesting that NuMA C-terminal fragment containing MTBD2 is also required to prevent hyper-clustering of NuMA. (D) Live fluorescent images of NuMA(N+C ΔNLS)-RFP-Nano (upper) and DHC-SNAP (lower). In 77% of cells (n=13), DHC was recruited to the cell cortex following light-induced targeting of NuMA (N+C ΔNLS)-RFP-Nano, but not in the remaining 23% of the cells. Scale bars = 10 μm.

**Figure S6:**
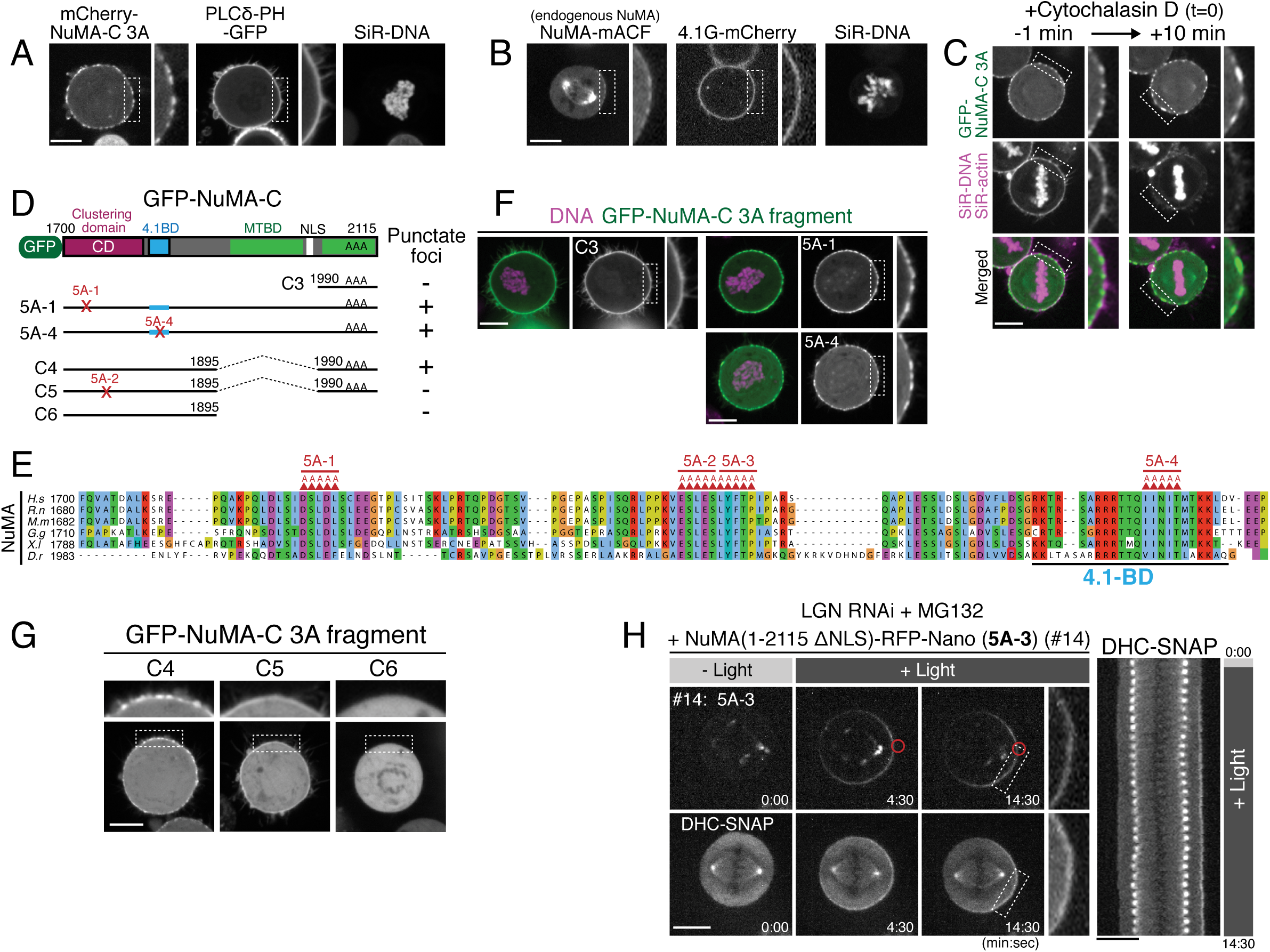
Identification of a clustering domain on NuMA’s C-terminal region. (A) Live fluorescence images of nocodazole-arrested HeLa cells showing mCherry-NuMA-C 3A and PLCΔ-PH-GFP, an indicator of PtdIns(4,5)P2 (Kotak et al., 2014). NuMA-C 3A, but not PLCΔ-PH, displays punctate signals. The localization of PLCΔ-PH was not affected by the expression of mCherry-NuMA-C 3A. DNA was visualized with SiR-DNA. (B) Live fluorescence images of endogenous NuMA-mACF and 4.1G-mCherry in HCT116 cells. NuMA-mACF, but not 4.1G-mCherry, shows punctate signals during prometaphase. (C) Live fluorescent images of HeLa cells showing GFP-NuMA-C 3A (upper) and SiR-DNA/SiR-actin (middle/left) following cytochalasin-D treatment (right). The NuMA foci preferentially accumulated at actin-poor cortical regions (left), and still localized following the disruption of actin polymerization (right). (D) GFP-tagged NuMA C-terminal fragment and the tested NuMA mutant fragments. 5A mutation sites are indicated in red. 3A mutation sites for CDK phosphorylation are shown in black AAA. (E) Amino acid sequence alignment of the 1700-1828 aa region of NuMA proteins aligned by ClustalWS. Highly conserved 5A mutation sites (5A-1 to 5A-4) are indicated by red triangles. (F) Live fluorescent images of nocodazole-arrested HeLa cells expressing GFP-tagged NuMA-C 3A fragments. NuMA C3 fragment (aa: 1990-2115) is sufficenet for cortical localization. NuMA-C 5A-1 and 5A-4 mutants still displayed punctate foci. (G) Live fluorescent images of nocodazole-arrested HeLa cells expressing GFP-tagged NuMA-C 3A fragments. Whereas NuMA(1700-1895: C6) diffused in the cytoplasm, this fragment was required for the NuMA (1990-2115: C3) fragment to display dots-like signals, which was abolished by 5A-2 mutation (C5). (H) Left: NuMA (1-2115 ΔNLS) 5A-3 (#10) mutant showing no cortical punctate signals of NuMA (top) and DHC-SNAP (bottom). Right: a kymograph obtained from image sequences of DHC-SNAP on the left at 30-sec intervals. The metaphase spindle was not fully displaced regardless of cortical dynein recruitment. Scale bars = 10 μm.

**Figure S7:**
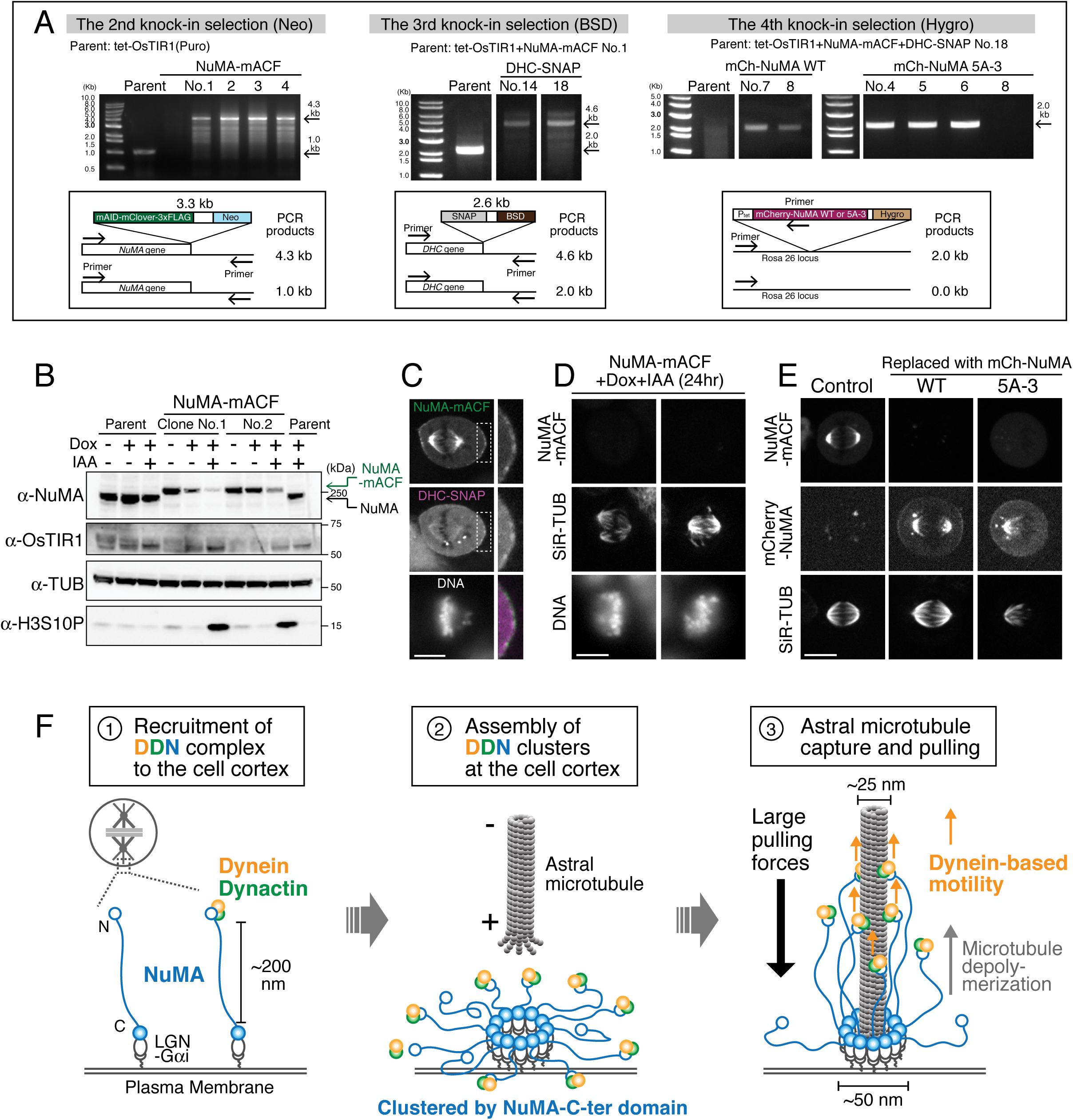
Auxin-inducible degradation of endogenous NuMA and its replacement with NuMA 5A-3 mutant. (A) Left: genomic PCR showing the genotype of clones after Neomycin (Neo) selection. All clones displayed a single 4.3-kb band, indicating that the mAID-mClover-3FLAG (mACF) (Neo) cassette was inserted at both *NuMA1* gene loci. The clone No.1 was used as a parent in the 3^rd^ selection. Middle: genomic PCR showing the genotype of clones after Blasticidin (BSD) selection. Both clones displayed a single 4.6-kb band, indicating that the SNAP (BSD) cassette was inserted in both *DHC1* gene loci. The clone No.18 was used as a parent in the 4^th^ selection. Right: genomic PCR showing the genotype of clones after Hygromycin (Hygro) selection. Arrows indicate a 2.0-kb band, which confirms the insertion of mCherry-NuMA WT or 5A-3 mutant cassette (Hygro) at the Rosa 26 locus. The clone WT (No.7) and 5A-3 (No.4) were used in this study. (B) Western blot probing for anti-NuMA, anti-OsTIR1, anti-α -tubulin (TUB, loading control), and anti-histone H3S10P (a mitotic marker) at 24 hr following treatment. Band shifts in NuMA-mACF indicate bi-allelic insertion of mAID tag. Treatment with both Dox and IAA caused degradation of NuMA-mACF and accumulation of phosphorylated Histone H3S10, indicating mitotic arrest or delay. (C) Live fluorescent images of endogenous NuMA-mACF and DHC-SNAP showing punctate signals at prometaphase cell cortex. These images were single z-sections captured with camera binning 1. DNA was visualized with Hoechst 33342. (D) Live fluorescent images of NuMA-mACF, SiR-tubulin (TUB) and DNA showing the spindle un-focusing phenotype. Intensity of SiR-TUB images was enhanced compared to Fig. 7B to improve clarity of the phenotypes. DNA was visualized with Hoechst 33342. (E) Live fluorescent images of endogenous NuMA-mACF, ectopically expressed mCherry-NuMA WT or 5A-3, and SiR-tubulin (TUB) showing spindle mis-orientation in the NuMA 5A-3 mutant cells. Single z-section images on the x-y plane are shown. (F) Model showing recruitment (#1) and assembly (#2) of the DDN cluster at the mitotic cell cortex, and multiple-arm capture and pulling of a single astral microtubule by the cortical DDN cluster (#3). See Discussion for details. Scale bars = 10 μm.

**Table S1:** Cell lines used in this study.

| No. | Name | Description | Clone No. | sgRNA+template plasmid | Parental cell | Reference |
| --- | --- | --- | --- | --- | --- | --- |
| 1 | HCT116 | HCT116 |  |  |  | ATCC CCL-247 |
| 2 | Mem-BFP-iLID | AAVS1::PCMV Mem-BFP-iLID (Puro) | 2 | AAVS1 T2 (Addgene#72833) and pTK511 | 1 | This study |
| 3 | NuMA-RFP-Nano | AAVS1::PCMV Mem-BFP-iLID (Puro), NuMA1::NuMA-tgRFPt-Nano (Neo) | 4 | pTK372 and pTK421 | 2 | This study |
| 4 | NuMA-RFP-Nano+DHC-SNAP | AAVS1::PCMV Mem-BFP-iLID (Puro), NuMA1::NuMA-tgRFPt-Nano (Neo), DHC1::DHC-SNAP (Hygro) | 8 | pTK308 and pTK565 | 3 | This study |
| 5 | NuMA-RFP-Nano+p150-SNAP | AAVS1::PCMV Mem-BFP-iLID (Puro), NuMA1::NuMA-tgRFPt-Nano (Neo), DCTN1::p150-SNAP (Hygro) | 15 | pTK525 and pTK587 | 3 | This study |
| 6 | Nano-mCherry-DHC | AAVS1::PCMV Mem-BFP-iLID (Puro), DHC1::Nano-mCherry-DHC (BSD) | 1 | pTK371 and pTK426 | 2 | This study |
| 7 | Nano-mCherry-DHC+p150-SNAP | AAVS1::PCMV Mem-BFP-iLID (Puro), DHC1::Nano-mCherry-DHC (BSD), DCTN1::p150-SNAP (Hygro) | 10 | pTK525 and pTK587 | 6 | This study |
| 8 | Nano-mCherry-DHC+NuMA-SNAP | AAVS1::PCMV Mem-BFP-iLID (Puro), DHC1::Nano-mCherry-DHC (BSD), NuMA1::NuMA-SNAP (Hygro) | 8 | pTK372 and pTK583 | 6 | This study |
| 9 | Mem-BFP-iLID+DHC-SNAP | AAVS1::PCMV Mem-BFP-iLID (Puro), DHC1::DHC-SNAP (BSD) | 11 | pTK308 and pTK600 | 2 | This study |
| 10 | DHC-SNAP+RFP-Nano | AAVS1::PCMV Mem-BFP-iLID (Puro), DHC1::DHC-SNAP (BSD), Rosa26::PTRE3G RFP-Nano (Hygro) | 18 | hROSA26 CRISPR-pX330 (Addgene#105927) and pTK630 | 9 | This study |
| 11 | DHC-SNAP+NuMA(1-2115)-RFP-Nano | AAVS1::PCMV Mem-BFP-iLID (Puro), DHC1::DHC-SNAP (BSD), Rosa26::PTRE3G NuMA(1-2115)-RFP-Nano (Hygro) | 10 | hROSA26 CRISPR-pX330 and pTK715 | 9 | This study |
| 12 | DHC-SNAP+NuMA(1-2115ΔNLS)-RFP-Nano | AAVS1::PCMV Mem-BFP-iLID (Puro), DHC1::DHC-SNAP (BSD), Rosa26::PTRE3G NuMA(1-2115ΔNLS)-RFP-Nano (Hygro) | 7 | hROSA26 CRISPR-pX330 and pTK713 | 9 | This study |
| 13 | DHC-SNAP+NuMA(1-1700)-RFP-Nano | AAVS1::PCMV Mem-BFP-iLID (Puro), DHC1::DHC-SNAP (BSD), Rosa26::PTRE3G NuMA(1-1700)-RFP-Nano (Hygro) | 28 | hROSA26 CRISPR-pX330 and pTK652 | 9 | This study |
| 14 | DHC-SNAP+NuMA(1-705)-RFP-Nano | AAVS1::PCMV Mem-BFP-iLID (Puro), DHC1::DHC-SNAP (BSD), Rosa26::PTRE3G | 5 | hROSA26 CRISPR-pX330 | 9 | This study |
|  |  | NuMA((1-705)-RFP-Nano (Hygro) |  | and pTK631 |  |  |
| 15 | DHC-SNAP+NuMA(1-705)4A-RFP-Nano | AAVS1::PCMV Mem-BFP-iLID (Puro), DHC1::DHC-SNAP (BSD), Rosa26:: PTRE3G NuMA(1-705)4A-RFP-Nano (Hygro) | 18 | hROSA26 CRISPR-pX330 and pTK660 | 9 | This study |
| 16 | DHC-SNAP+NuMA(214-705)-RFP-Nano | AAVS1::PCMV Mem-BFP-iLID (Puro), DHC1::DHC-SNAP (BSD), Rosa26:: PTRE3G NuMA(214-705)-RFP-Nano (Hygro) | 2 | hROSA26 CRISPR-pX330 and pTK643 | 9 | This study |
| 17 | DHC-SNAP+NuMA(1-505)-RFP-Nano | AAVS1::PCMV Mem-BFP-iLID (Puro), DHC1::DHC-SNAP (BSD), Rosa26:: PTRE3G NuMA(1-505)-RFP-Nano (Hygro) | 21 | hROSA26 CRISPR-pX330 and pTK664 | 9 | This study |
| 18 | DHC-SNAP+NuMA(1-413)-RFP-Nano | AAVS1::PCMV Mem-BFP-iLID (Puro), DHC1::DHC-SNAP (BSD), Rosa26:: PTRE3G NuMA(1-413)-RFP-Nano (Hygro) | 23 | hROSA26 CRISPR-pX330 and pTK641 | 9 | This study |
| 19 | DHC-SNAP+NuMA(1-213)-RFP-Nano | AAVS1::PCMV Mem-BFP-iLID (Puro), DHC1::DHC-SNAP (BSD), Rosa26:: PTRE3G NuMA(1-213)-RFP-Nano (Hygro) | 8 | hROSA26 CRISPR-pX330 and pTK642 | 9 | This study |
| 20 | DHC-SNAP+NuMA(1-1895)-RFP-Nano | AAVS1::PCMV Mem-BFP-iLID (Puro), DHC1::DHC-SNAP (BSD), Rosa26:: PTRE3G NuMA(1-1895)-RFP-Nano (Hygro) | 4 | hROSA26 CRISPR-pX330 and pTK678 | 9 | This study |
| 21 | DHC-SNAP+NuMA(1-1985)-RFP-Nano | AAVS1::PCMV Mem-BFP-iLID (Puro), DHC1::DHC-SNAP (BSD), Rosa26:: PTRE3G NuMA(1-1985)-RFP-Nano (Hygro) | 19 | hROSA26 CRISPR-pX330 and pTK711 | 9 | This study |
| 22 | DHC-SNAP+NuMA(1-2115Δex24)-RFP-Nano | AAVS1::PCMV Mem-BFP-iLID (Puro), DHC1::DHC-SNAP (BSD), Rosa26:: PTRE3G NuMA(1-2115Δex24)-RFP-Nano (Hygro) | 17 | hROSA26 CRISPR-pX330 and pTK728 | 9 | This study |
| 23 | DHC-SNAP+NuMA(1-705+1700-2115ΔNLS)-RFP-Nano | AAVS1::PCMV Mem-BFP-iLID (Puro), DHC1::DHC-SNAP (BSD), Rosa26:: PTRE3G NuMA(1-705+1700-2115ΔNLS)-RFP-Nano (Hygro) | 4 | hROSA26 CRISPR-pX330 and pTK729 | 9 | This study |
| 24 | DHC-SNAP+NuMA(1700-2115)-RFP-Nano | AAVS1::PCMV Mem-BFP-iLID (Puro), DHC1::DHC-SNAP (BSD), Rosa26:: PTRE3G NuMA(1700-2115)-RFP-Nano (Hygro) | 5 | hROSA26 CRISPR-pX330 and pTK645 | 9 | This study |
| 25 | DHC-SNAP+NuMA(1-2115ΔNLS)5A-3-RFP-Nano | AAVS1::PCMV Mem-BFP-iLID (Puro), DHC1::DHC-SNAP (BSD), Rosa26:: PTRE3G NuMA(1-2115ΔNLS)5A-3-RFP-Nano (Hygro) | 4 | hROSA26 CRISPR-pX330 and pTK746 | 9 | This study |
| 26 | HCT116 tet-OsTIR1 | AAVS1::PTRE3G OsTIR1 (Puro) |  | pAAVS1 T2 and MK243 (Addgene#72835) | 1 | (Natsume et al., 2016) |
| 27 | NuMA-mACF | AAVS1::PTRE3G OsTIR1 (Puro), NuMA1:: NuMA-mAID-mClover-3FLAG (Neo) | 1 | pTK372 and pTK398 | 26 | This study |
| 28 | NuMA-mACF+DHC-SNAP | AAVS1::PTRE3G OsTIR1 (Puro), NuMA1:: NuMA-mAID-mClover-3FLAG (Neo), DHC1:: DHC-SNAP (BSD) | 18 | pTK308 and pTK600 | 27 | This study |
| 29 | NuMA-mACF+DHC-SNAP+mCh-NuMA WT | AAVS1::PTRE3G OsTIR1 (Puro), NuMA1:: NuMA-mAID-mClover-3FLAG (Neo), DHC1:: DHC-SNAP (BSD), Rosa26:: PTRE3G mCherry-NuMA(WT) (Hygro) | 7 | hROSA26 CRISPR-pX330 and pTK503 | 28 | This study |
| 30 | NuMA-mACF+DHC-SNAP+mCh-NuMA 5A-3 | AAVS1::PTRE3G OsTIR1 (Puro), NuMA1:: NuMA-mAID-mClover-3FLAG (Neo), DHC1:: DHC-SNAP (BSD), Rosa26:: PTRE3G mCherry-NuMA(5A-3) (Hygro) | 4 | hROSA26 CRISPR-pX330 and pTK750 | 28 | This study |
| 31 | HeLa | HeLa |  |  |  | (Kiyomitsu & Cheeseman, 2012) |

**Table S2:**
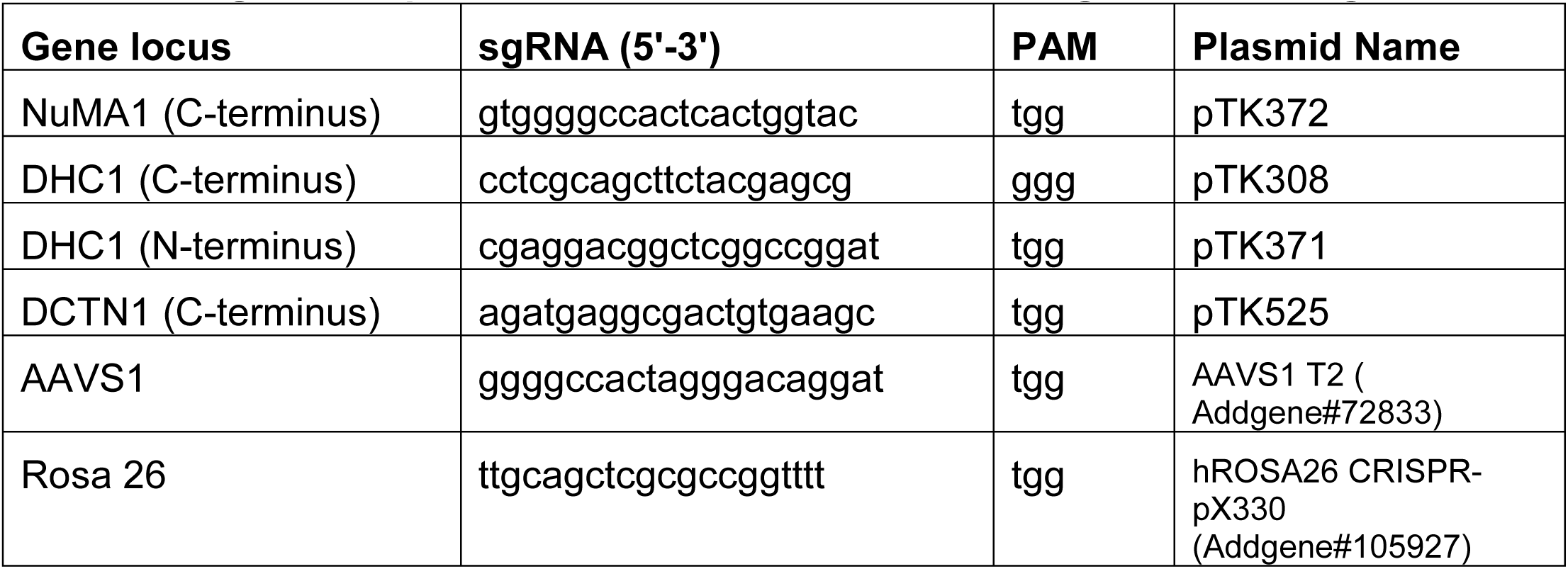
sgRNA sequences for CRISPR/Cas9-mediated genome editing

**Table S3:**
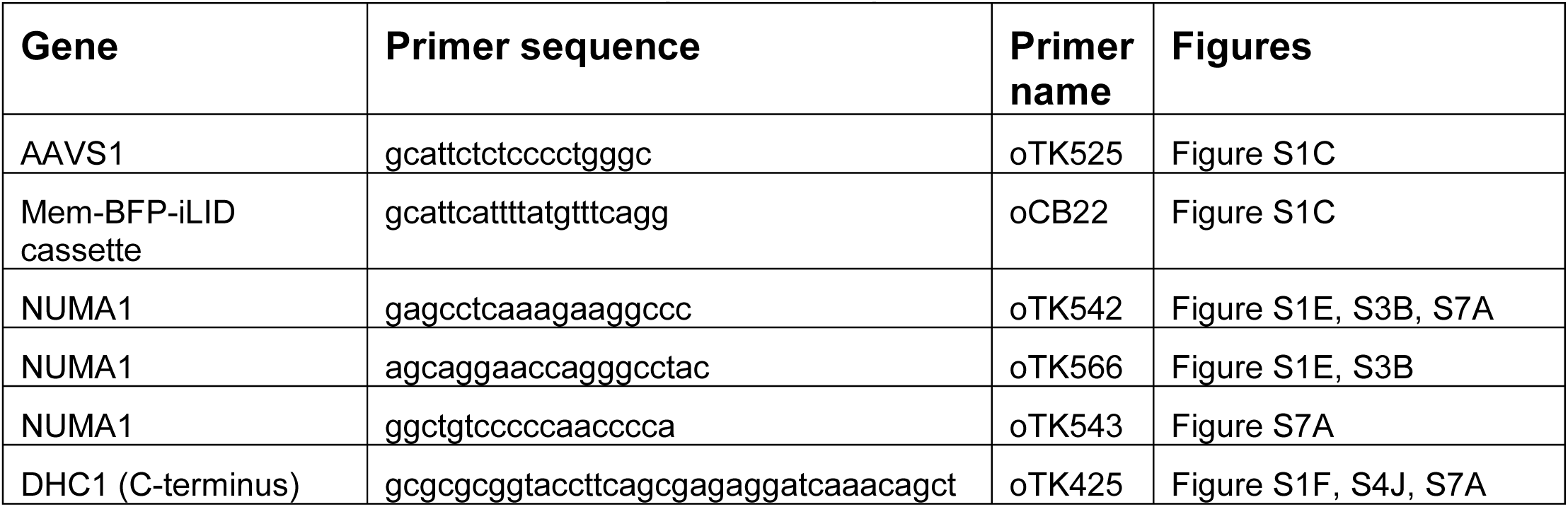
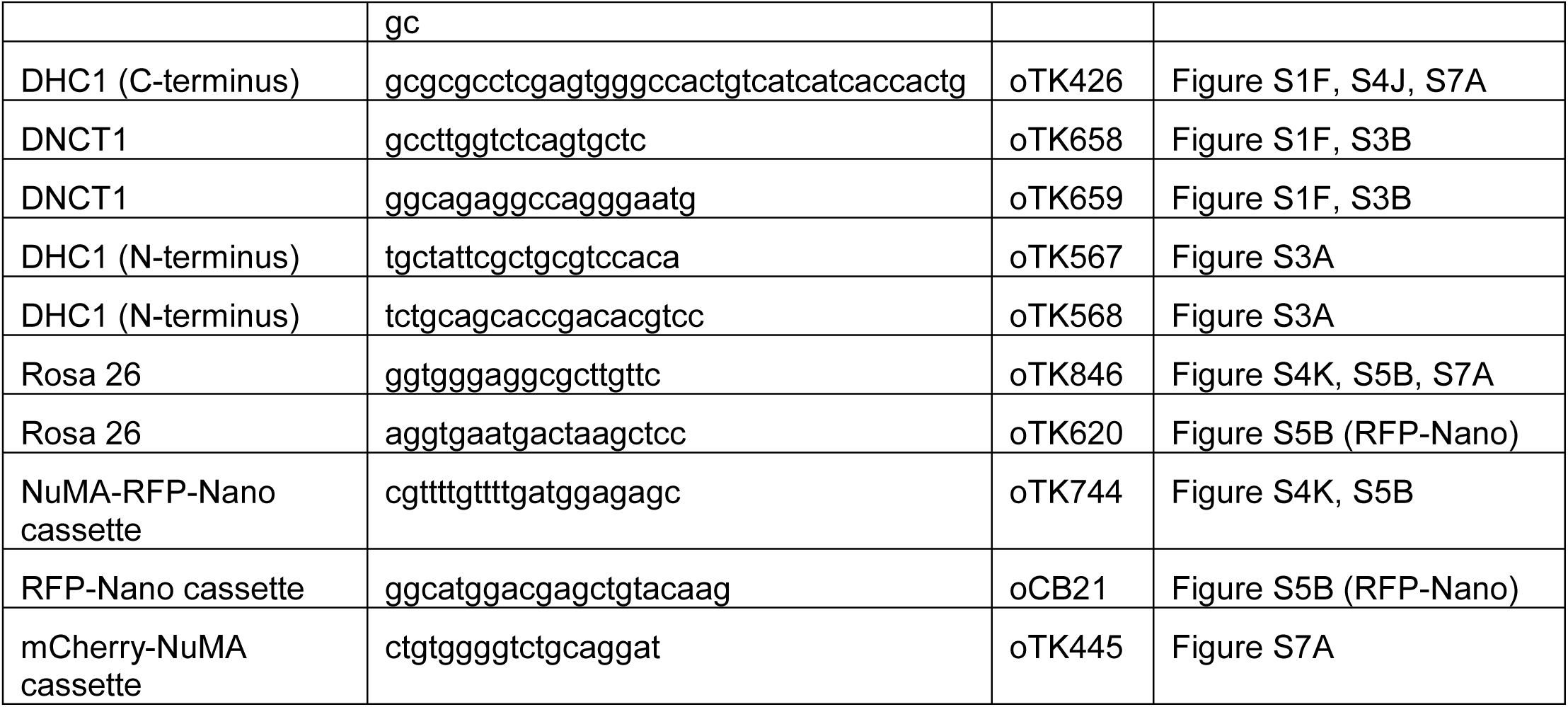
PCR primers to confirm gene editing

**Movie S1: Light-induced cortical targeting of RFP-Nano.** The dynamic cortical targeting and repositioning of RFP-Nano, in response to illuminations, are shown in this movie; it is played at 5 fps.

**Movie S2: Light-induced cortical targeting of NuMA-RFP-Nano and spindle pulling.** Light-induced cortical recruitment of NuMA-RFP-Nano (left), and DHC-SNAP (right), and spindle displacement toward NuMA/DHC-enriched cell cortex have been shown in this movie; it is played at 5 fps.

**Movie S3: Light-induced cortical repositioning of NuMA-RFP-Nano and spindle rotation.** Light-induced cortical repositioning of NuMA-RFP-Nano (left), and dynamics of SiR-tubulin (right) have been shown in this movie. The spindle rotation was coupled with cortical repositioning of NuMA. This movie is played at 5 fps.

